# Analyzing asymmetry in brain hierarchies with a linear state-space model of resting-state fMRI data

**DOI:** 10.1101/2023.11.04.565625

**Authors:** Danilo Benozzo, Giacomo Baggio, Giorgia Baron, Alessandro Chiuso, Sandro Zampieri, Alessandra Bertoldo

## Abstract

The study of functional brain connectivity in resting-state functional magnetic resonance imaging (rsfMRI) data has traditionally focused on zero-lag statistics. However, recent research has emphasized the need to account for dynamic aspects due to the complex patterns of time-varying co-activations among brain regions. In this regard, the importance of non-zero-lag statistics in studying complex brain interactions has been emphasized, both in terms of modeling and data analysis. Here, we show how a time-lag description is incorporated within the framework of dynamic causal modeling (DCM) resulting in an asymmetric state interaction matrix known as effective connectivity (EC). Asymmetry in EC is conventionally associated with the directionality of interactions between brain regions and is frequently employed to distinguish between incoming and outgoing node connections. We will revisit this interpretation by employing a decomposition of the EC matrix. This decomposition enables us to isolate the steady-state differential cross-covariance matrix, which is responsible for modeling the information flow and introducing time irreversibility. In other words, by modeling the off-diagonal part of the differential covariance, the system landscape may exhibit a curl steady-state flow component that breaks detailed balance and diverges the dynamics from equilibrium. Our empirical results reveal that only the outgoing strengths of the EC matrix relate with the flow described by the differential cross-covariance, while the so-called incoming strengths are primarily driven by the zero-lag covariance, specifically the precision matrix, thus reflecting conditional independence rather than directionality.

## 1 Introduction

An excellent framework for comprehending how the brain spontaneously orchestrates its activities is given by whole-brain resting-state recordings acquired through functional magnetic resonance imaging (fMRI). After substantial efforts aimed at characterizing functional connectivity (FC), which measures steady-state similarity between pairs of brain regions, there has been an increasing focus on investigating dynamic properties [41, 69, 37]. Multiple perspectives have been utilized to provide insights into this domain. These range from data-driven approaches, which seek to describe the time variability of functional connectivity (referred to as dynamical FC or dFC) [1, 59, 36], to more complex dynamical systems models that aim to replicate the dynamic structure of the state space [7, 78].

Dynamic causal modelling (DCM) is one of the prominent methods used for modeling fMRI data in a top-down approach [28]. Initially, DCM was conceived to model task-based recordings encompassing a restricted number of regions, within a framework of competing hypotheses. However, more recently, DCM has been extended to encompass whole-brain recordings obtained during resting-state conditions [31, 62]. In any scenario, important aspects of DCM include its Bayesian inference framework, the state-space model with linear state equation, and the non-linear observational equation designed to address the relationship between neural activity and BOLD signal due to hemodynamic coupling [30]. Of significance within this context is the state interaction matrix, referred to as effective connectivity (EC), which captures the causal impact that each component of the system has on the behavior of the other components [9]. The linearity of the state space confines the reconstruction of the phase space around an isolated fixed point, resulting in a first-order approximation of the underlying vector field. This property makes the effective connectivity (EC) to correspond to the Jacobian matrix. On one side, this poses a limitation that makes it challenging to fully grasp non-linear phenomena like metastability and phase transitions [16]. However, on the flip side, single-fixed point dynamical models have demonstrated their effectiveness in describing large-scale brain activity [56, 75, 74]. And this serves as a trade-off, rendering an ill-posed problem more manageable.

Here, we focus on the asymmetry of EC, which is an indicator of the non-equilibrium steady-state (NESS) regime [27], along with its implications. This involves a curl steady-state flow in the system landscape that is connected to a non-zero entropy production rate, thus underlining the dynamics’ time-irreversibility [77]. Specifically, the notion of directionality emerges inherently from the direction of rotation of the steady-state flow which is decoupled in its solenoidal component, enabling the recognition of brain regions that act as senders or receivers, thus forming a directed hierarchy. Under our modelling hypothesis, the solenoidal component is driven by the off-diagonal part of the differential covariance, which we will refer to as the differential cross-covariance (dC-Cov). The differential covariance is essentially a covariance matrix that quantifies how one node influences the time derivative of another [48, 49].

We consider it important to shed light on this topic for the following reasons: i) EC has frequently been portrayed as a directed counterpart of FC in a graph-based manner, often overlooking its underlying context in dynamical system theory. ii) Considering the substantial focus on investigating cortical functional hierarchies at the macro-scale [54, 42], the ability to pinpoint the direction through which information traverses such hierarchies would significantly enhance our comprehension of the phenomenon. iii) Despite potential limitations, the linear approximation offers a distinct advantage by enabling the closed-form calculation of the curl steady-state flow, and related metrics, such as the entropy production rate [81, 32].

This work aims to show how the asymmetry of EC, which is inherently linked to a non-zero dC-Cov, is reflected in the neuronal state in terms of power spectral density and time-irreversibility. This will provide a mechanistic justification for the presence of ultraslow fluctuations, which are a characteristic phenomenon in resting-state recordings [51].

The final part of our work will concentrate on interpreting the row and column strengths of the EC matrix. These are commonly used measures to quantify the incoming and outgoing connectivity of each node [64, 33]. We will consider that EC can be decomposed into the product of the differential covariance and the precision matrices. Meaning that each row of the differential covariance is stretched toward the principal components of the precision matrix. By this decomposition into zero-lag covariance (the precision matrix) and differential covariance, we will derive how they contribute to defining the row and column strengths of EC. Our empirical findings revealed that EC row strengths are directly related to the corresponding node strengths of the precision matrix, reflecting conditional independence rather than indicating directionality. On the other hand, the column sums of EC are associated with the corresponding column sums of the steady-state dC-Cov, offering insights into directionality. Since this latter shapes the solenoidal steady-state flow which determines the direction of information propagation, our empirical findings suggest that only the column strengths of EC convey a directional interpretation. Specifically, a large positive column sum suggests a dominant source/sender behavior of the node, while a large negative column sum indicates a dominant sink/receiver behavior. We conclude by providing three examples of applying dC-Cov to uncover the underlying directed hierarchy in resting-state fMRI data, using a mouse dataset and two distinct human datasets.

## 2 Materials and Methods

### 2.1 Dataset

#### 2.1.1 Mouse dataset

A dataset of n=20 adult male C57BI6/J mice were previously acquired at the IIT laboratory (Italy). All in vivo experiments were conducted in accordance with the Italian law (DL 2006/2014, EU 63/2010, Ministero della Sanità, Roma) and the recommendations in the Guide for the Care and Use of Laboratory Animals of the National Institutes of Health. Animal research protocols were reviewed and consented by the animal care committee of the Italian Institute of Technology and Italian Ministry of Health. Animal preparation, image data acquisition and image data preprocessing for rsfMRI data have been described in greater detail elsewhere [38]. Briefly, rsfMRI data were acquired on a 7.0-T scanner (Bruker BioSpin, Ettlingen) equipped with BGA-9 gradient set, using a 72 mm birdcage transmit coil, and a four-channel solenoid coil for signal reception. Single-shot BOLD echo planar imaging time series were acquired using an echo planar imaging sequence with the following parameters: repetition time/echo time, 1000/15 ms; flip angle, 30°; matrix, 100×100; field of view, 2.3 ×2.3 cm^2^; 18 coronal slices; slice thickness, 0.60 mm; 1920 volumes.

Regarding image preprocessing as described in [39], timeseries were despiked, motion corrected, skull stripped and spatially registered to an in-house EPI-based mouse brain template. Denoising and motion correction strategies involved the regression of mean ventricular signal plus 6 motion parameters [65]. The resulting timeseries were then band-pass filtered (0.01-0.1 Hz band). After preprocessing, mean regional time-series were extracted for 74 (37+37) regions of interest (ROIs) derived from a predefined anatomical parcellation of the Allen Brain Institute (ABI), [57, 79].

#### 2.1.2 MPI-LMBB dataset

A subset of 295 subjects has been selected from the resting-state scans of the publicly available MPI-Leipzig Mind-Brain-Body dataset (https://legacy.openfmri.org/dataset/ds000221/). The data selection was performed on the original dataset (consisting of 318 individuals) by excluding participants with high motion or pre-precessing failures. Then, half of the dataset was utilized for clustering purposes (detailed in the following section). From the remaining portion, we selected only younger participants aged between 20 and 30 years, resulting in a final sample of 107 subjects.

Data acquisition was performed with a 3T Siemens Magnetom Verio scanner, equipped with a 32-channel head coil. The protocol included a T1-weighted 3D magnetization-prepared 2 rapid acquisition gradient echoes (3D-MP2RAGE), and resting-state fMRI images acquired using a 2D multiband gradient-recalled echo echo-planar imaging (EPI) sequence (TR = 1400 ms; flip angle = 69°; voxel size = 2.3×2.3×2.3mm; volumes = 657; multiband factor = 4) and two spin echo acquisitions. For additional information regarding the acquisition protocol, please refer to [3]. Details concerning the preprocessing of rsfMRI data can be found in [73]. Subsequently, the temporal traces were subjected to band-pass filtering in the frequency range of 0.0078 to 0.1 Hz and temporally despiked by means of the *icatb_despike_tc* function of the GIFT toolbox (http://trendscenter.org/software/gift/).

#### 2.1.3 HCP Young Adult dataset

We used the HCP Yuong Adult (HCP-YA) data release, which included 1,200 normal young adults, aged 22-35 [26]. All data were collected on a 3T Siemens Skyra scanner with gradients customized for the HCP. We restricted our analysis to 173 subjects (age range 22-35 yo). rfMRI data were acquired in four runs of approximately 15 minutes each, two runs in one session and two in another session, with eyes open with relaxed fixation (sequence parameters for each run: gradient-echo EPI, TR = 720 ms, TE = 33.1 ms, flip angle = 52°, FOV = 208×180, voxel size = 2 mm isotropic, frames=1200). Within each session, oblique axial acquisitions alternated between phase encoding in a right-to-left (RL) direction in one run and phase encoding in a left-to-right (LR) direction in the other run [26].

Concerning data preprocessing, the dataset used in this study was the preprocessed volumetric version provided by HCP [34] which combines a set of tools from FSL, FreeSurfer, and the HCP Connectome Workbench [53]. It encompasses three MR structural pipelines for distortion correction and the alignment of individual brain data to a standardized MNI template. The fMRI data were preprocessed using the MR functional pipeline, which involved distortion correction, motion correction by aligning fMRI volumes, and registration of the fMRI data to the corresponding structural data. For each subject, only the second run (run2) of the data was utilized. Additionally, the data were temporally filtered with a band-pass range of 0.0078-0.1 Hz and temporally despiked by means of the *icatb_despike_tc* function of the GIFT toolbox.

### 2.2 Cortical network partitions

In the mouse data, we partitioned the cortical areas into six macro-regions [40]. These regions include: the somatomotor area (SM) which corresponds to the functional lateral cortical network [50], the visual area (Vis), the temporal area (TEMP), the anterolateral area (antL) which mainly overlaps with the salience network (agranular insula areas) and part of the default mode posterolater network (DMN-postl), the medial area (MED) representing part of the default mode midline network (DMNmid) [39] and the prefrontal area (PF), which completes the anterior part of DMNmid. In terms of subcortical regions, our adopted parcellation includes: the hippocampus (HP), the striatum regions (STR), the basal forebrain (comprising the lateral septal complex, striatum-like amygdalar nuclei and pallidum), the thalamus and the hypothalamus. For a comprehensive list, please refer to Supplementary Table S1. Regarding the human datasets, in both cases we extracted time series data from a 100-area parcellation scheme of the cortex provided by the Schaefer atlas [68], which maps to 7 resting-state functional networks (RSNs) referring to both hemispheres: Visual network (Vis) (5 parcels), Somatomotor network (SomMot) (6 parcels), Dorsal attention network (DorsAttn) (9 parcels), Saliency/Ventral attention network (SalVenAttn) (11 parcels), Limbic network (Limbic) (5 parcels), Control network (Cont) (10 parcels), Default mode network (DMN/Default) (16 parcels). We also defined a set of 12 subcortical regions based on the AAL2 segmentation [66]. For each hemisphere, we selected 6 regions consisting of thalamus proper, caudate, putamen, pallidum, hippocampus and cerebellum.

A consensus clustering procedure was applied to reduce the number of cortical parcels to reach a parcellation compatible with the DCM inference algorithm. As a result, we applied to the first half of the MPI-LMBB dataset (147 subjects) a Consensus Clustering Evidence Accumulation (CCEA) procedure, similarly to what described in [67], to determine the number of optimal clusters. Specifically, this framework employs base and consensus clustering methods to get robust and stable clusters. Details on the procedure can be found in [4]. The clustering procedure yielded 62 cortical clusters that, when combined with the 12 subcortical regions, resulted in a total of 74 parcels. For a comprehensive list, please refer to Supplementary Table S2.

### 2.3 State-space linear model

The linear DCM framework employs a linear stochastic model to describe the neuronal state. This model represents a continuous-time linear time-invariant stochastic system

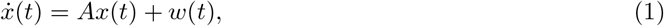

where *x* ∈ ℝ ^*n*^ is the vector containing the states of the network nodes, *A* ∈ ℝ^*n×n*^ is the state interaction matrix (the so-called effective connectivity matrix, EC), and the input *w* is a zero-mean white noise vector with positive definite covariance matrix Σ_*w*_ = 𝔼 [*w*(*t*)*w*(*t*)^*T*^]. We assumed uncorrelated noise across nodes so that Σ_*w*_ = *σ* ^2^*I*_*n*_ where *σ* ^2^ represents the variance of each brain endogenous fluctuation and *I*_*n*_ the identity matrix of size *n*.

Moreover, *A* is assumed to be stable (its all eigenvalues have strictly negative real part), this ensures the existence of a positively definite steady-state covariance matrix Σ, which, after normalization, represents the functional connectivity matrix (FC)

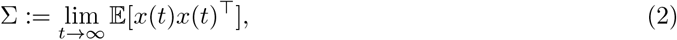

for which the algebraic Lyapunov equation holds

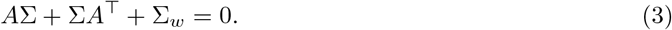

Importantly, *A* can be decomposed such that

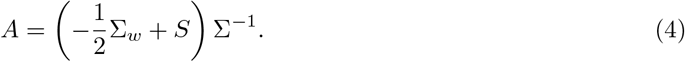

Eq. 4 parametrizes all *A* matrices that give the same Σ, i.e. the covariance matrix of the state, for any *S* skew-symmetric matrix, i.e. *S* = *−S*^*T*^. This defines a one-to-one map between each (Σ, *S*) and *A* given the noise covariance Σ_*w*_ [12], with 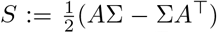. Any off-diagonal element (*i, j*) of *S* can be written in statistical term as 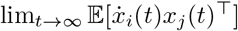, and we refer to this matrix as the steady-state differential cross-covariance (dC-Cov). Similarly, the differential auto-covariance (dA-Cov) is represented by −Σ_*w*_*/*2. The sum of these two matrices constitutes the differential covariance matrix [49].

Eq. 4 can also be seen as following from the fundamental theorem of vector calculus (or Helmholtz decomposition), which states that any vector field in steady state can be broken down into the combination of a dissipative (curl-free) flow which is mediated by −Σ_*w*_*/*2 (dA-Cov), and a solenoidal (divergence-free) flow which is mediated by *S* (dC-Cov) [82, 27]. The existence of both these flows constitutes the core of non-equilibrium steady-state dynamics. A non-zero *S* matrix introduces time irreversibility into the dynamics, and inverting the sign of *S* results in the reversal of dynamics [18].

The differential cross-covariance matrix *S* gives a direct interpretation of which node behaves as source and which as sink [49, 15]. If *S*_*i,j*_ *>* 0 then node *j* is considered the source, and node *i* is considered the sink, indicating that information is flowing from node *j* to node *i*. In the context of *S* being skew-symmetric, this example implies *S*_*j,i*_ *<* 0. Therefore, to maintain coherence with the established interpretation, the row entry corresponds to the source and the column entry corresponds to the sink.

This interpretation provides a straightforward method to quantify the prevalence of a node acting as a source or sink within the system. For example, summing the values in each column of the *S* matrix allows us to gauge the extent to which a node predominantly serves as a source (if the sum is largely positive), a sink (if the sum is largely negative), or maintains balance between the two roles (if the sum is close to zero). A sum of zero may also indicate that the node is excluded from contributing to the steady-state solenoidal flow.

In characterizing nodes within a network, matrix *A*, representing effective connectivity, is commonly employed to quantify incoming and outgoing strengths. Typically, for a given node, its incoming and outgoing strengths are calculated as the sums of the node’s corresponding row and column in matrix *A*, respectively. In light of our previous part about the pivotal role of *S* in shaping the directionality of steady-state dynamics, our objective is to study how this is reflected on the asymmetry of *A*. In the following, we will use the notation *A*_*i*,._ and *A*_.,*i*_ to represents the row and column sum of node *i* in *A*, respectively. If *A* is symmetric the following holds:

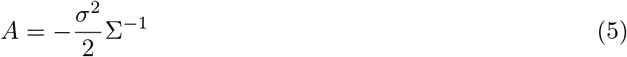

Thus, *A* is the outcome of the multiplication of the dA-Cov by the precision matrix, yielding a scaled version of the precision matrix. Since the precision matrix is symmetric and the noise variance is assumed equal across regions, incoming and outgoing strengths of any node *i* are equal:

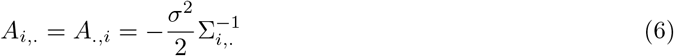

While in the presence of causal interactions among regions, which corresponds to a non-zero dC-Cov, i.e. *S ≠* 0, from Eq. 4 it follows that the incoming strength of node *i* becomes:

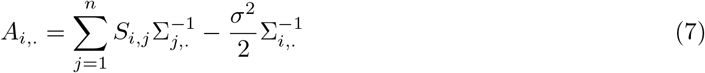

that is a linear combination computed by taking the sum of rows (or columns) of the precision matrix, i.e. 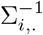, where each row sum (or column sum) is weighted by the corresponding element in the *i*-th row of matrix *S*, plus the term in Eq. 6. Similarly, the outgoing strength of node *i* becomes:

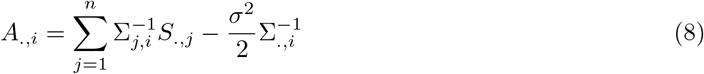

thus a linear combination where each *S* row sum is weighted by the corresponding element in the *i*-th column of the precision matrix, plus 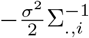, i.e. the strength in the symmetric case Eq. 6.

### 2.4 sparse-DCM

The effective connectivity matrix, along with other model variables, was estimated at the single-subject level using the method described in [62], which is referred to as sparse-DCM. In line with the DCM framework [28], sparse-DCM is a state-space model where the state *x*(*t*) satisfies a set of linear differential equations representing the coupling among neural components, and the output model maps the neuronal activity to the measured BOLD signal *y*(*t*) through the hemodynamic response function (HRF):

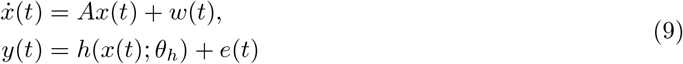

with *A* representing the effective connectivity matrix, *h*(.) the hemodynamic response that is modeled by the biophysically inspired Balloon-Windkessel model [30] and *θ*_*h*_ its parameters. *w*(*t*) denotes the stochastic intrinsic brain fluctuations and *e*(*t*) the observation noise, both are Gaussian variables with zero mean and diagonal covariance matrices *σ*^2^*I*_*n*_ and *R* = diag(*λ*_1_, *λ*_2_, …, *λ*_*n*_), respectively.

To address the computational burden of model inversion when dealing with whole brain data, [62] proposed a discretization and linearization of Eq. 9 as well as a sparsity-inducing prion on the EC matrix, i.e *A* in Eq. 9. This was motivated by the low temporal resolution of fMRI data, which usually ranges from 0.5 to 3 seconds, and the idea that the hemodynamic response *h*(.) can be modeled as a Finite Impulse Response (FIR) model with input the neuronal state and output the BOLD signal. In our study, to ensure that the length of the input response was large enough to model relevant temporal dependencies, we set the hemodynamic length to 18 samples with a sampling time 1s (TR). For each brain parcel *i*, a finite impulse response *h*_*i*_ ∼ *𝒩* (*μ*_*h*_, Σ_*h*_) was assigned by deriving *μ*_*h*_ and Σ_*h*_ through a Monte-Carlo sampling of typical responses generated by the non-linear Balloon-Windkessel model (10000 samples). The sparsity-inducing prior on the EC estimation was formulated to reduce as much as possible spurious couplings. In particular, each element *a*_*i*_ of matrix *A* was assumed to be a Gaussian variable with zero mean and *γ*_*i*_ variance. The hyperparameter *γ* = [*γ*_1_, *γ*_2_, …, *γ*_*n*_^2^] was estimated through marginal likelihood maximization. Under generic conditions, the maximum likelihood estimate of some *γ*_*i*_-s will be zero such that the Gaussian posterior distribution of their corresponding *a*_*i*_ is concentrated around zero thus producing a zero MAP estimate. In sparse-DCM, model inversion and parameter optimization are performed by an expectation-maximization (EM) algorithm.

### 2.5 Metric description

#### 2.5.1 Metrics of time-irreversal asymmetry

From [32], the entropy production rate of a linear stochastic model is

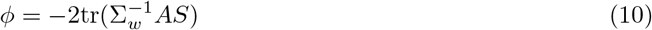

which is a scale measure that quantifies the degree of irreversibility of the whole process. The local contribution of each node to the global entropy production rate is quantified through the nodeirreversibility as Σ_*i*_ |*S*_*ij*_|. Both metrics are zero in reversible dynamics, indicating no entropy production, while they are greater than zero in irreversible non-equilibrium dynamics.

With the chosen linear state-space model, a non-zero *S* matrix directly influences the distribution of energy within the neuronal state across different frequencies, i.e. the power spectral density Φ(*ιω*) [12]:

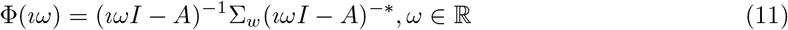

with denoting the conjugate transpose. In particular, the power spectral density of each neuronal state, at a specific frequency *ω*, corresponds to the related diagonal element in Φ(*ιω*), while the offdiagonal elements represent the cross-spectral densities. In the case of time-reversible dynamics, it is expected that the power spectral density will decrease as the frequency *ω* increases. Conversely, for non-time-reversible dynamics, it is anticipated to peak at a frequency greater than 0 [12].

#### 2.5.2 Metastability and synchronization indices

Metastability and synchronization are indices that describe the phase coherence of a signal ensemble over time [10]. The phase coherence of *n* signals is evaluated at each time bin using the Kuramoto order parameter

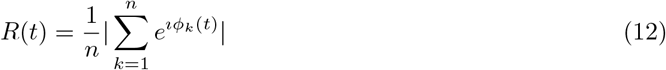

where *Φ*_*k*_ (*t*) represents the instantaneous phase of the fMRI signal for node *k*. The value of *R*(*t*) ranges from 0 to 1, where 1 indicates full synchronization and 0 represents randomness - phases uniformly distributed between 0 and 2*π*. The synchronization index is calculated as the mean of *R*(*t*) over time, while the metastability index represents its standard deviation [71]. The instantaneous phase *Φ*_*k*_ (*t*) of *x*_*k*_ (*t*) was computed as the argument of the complex signal *z*_*k*_ (*t*) = *x*_*k*_ (*t*) + *iH*[*x*_*k*_ (*t*)], with *H*[.] denoting the Hilbert transformation. By denoting the module of *z*_*k*_ (*t*) with *M*_*k*_ (*t*), *x*_*k*_ (*t*) can be represented as a rotating vector with phase *Φ*_*k*_ (*t*) and magnitude *M*_*k*_ (*t*), i.e. *x*_*k*_ (*t*) = *M*_*k*_ (*t*) cos *Φ*_*k*_ (*t*).

#### 2.5.3 DCM as generative model

After fitting a DCM model to each empirical single-subject rsfMRI recording, we utilized the model to generate synthetic realizations of rsfMRI signals. This procedure was carried out for each dataset using two different approaches: the standard implementation of sparse-DCM proposed in [62], and a constrained version that enforced the state-space matrix *A* to be symmetric, thereby imposing time-reversibility on the dynamics.

Synthetic data were employed to assess the model’s ability to generate data exhibiting static and dynamic functional connectivity (sFC and dFC, respectively) that resembles the empirical counterparts. To quantify the similarity between sFC, we computed the correlation between the triangular part of empirical and simulated sFC. For dFC, the similarity was determined using the Kolmogorov-Smirnov distance between the triangular parts of empirical and simulated dFC. The calculation of dFC was performed using a sliding window of 50 s with a step of 25 s.

Synthetic data were also used to assess how the synchronization and metastability indices change in data generated by both time-irreversible and time-reversible systems. We aimed to evaluate and compare these indices between the synthetic data and the empirical data, thus gaining insights into the differences arising from the two different dynamics.

## 3 Results

We applied sparse-DCM at the single-subject level, and identified the related model. We employed both the model and the implementation described in [62]. Initially, we calculated the dC-Cov (*S* matrix) and focused on its influence on both the power spectra density and its time-domain counterpart, i.e. the auto-correlation, of the neuronal state.

A non-zero dC-Cov has a direct impact on the power spectral density of the neuronal state, causing a shift in the frequency peak towards values greater than zero, as depicted in Fig. 1b. In other words, a non-zero node irreversibility corresponds to a frequency peak greater than zero in the psd, as shown in Fig. 1c. Similarly, in the time-domain, the auto-correlation exhibits a negative peak when non-zero node irreversibility is present, whereas it decreases monotonically, as shown in Fig. 1d. When *S* is constrained to be zero by either deriving a new *A* matrix enforcing *S* = 0 from Eq. 4 (refer to Fig. 1e) or by identifying a time-reversible model (see Fig. 1f), the auto-correlation is consistently positive and decreases with frequency. The imposition of time-reversibility results in increasing the memory of the system due to a slower decay in auto-correlation.

**Figure 1.**
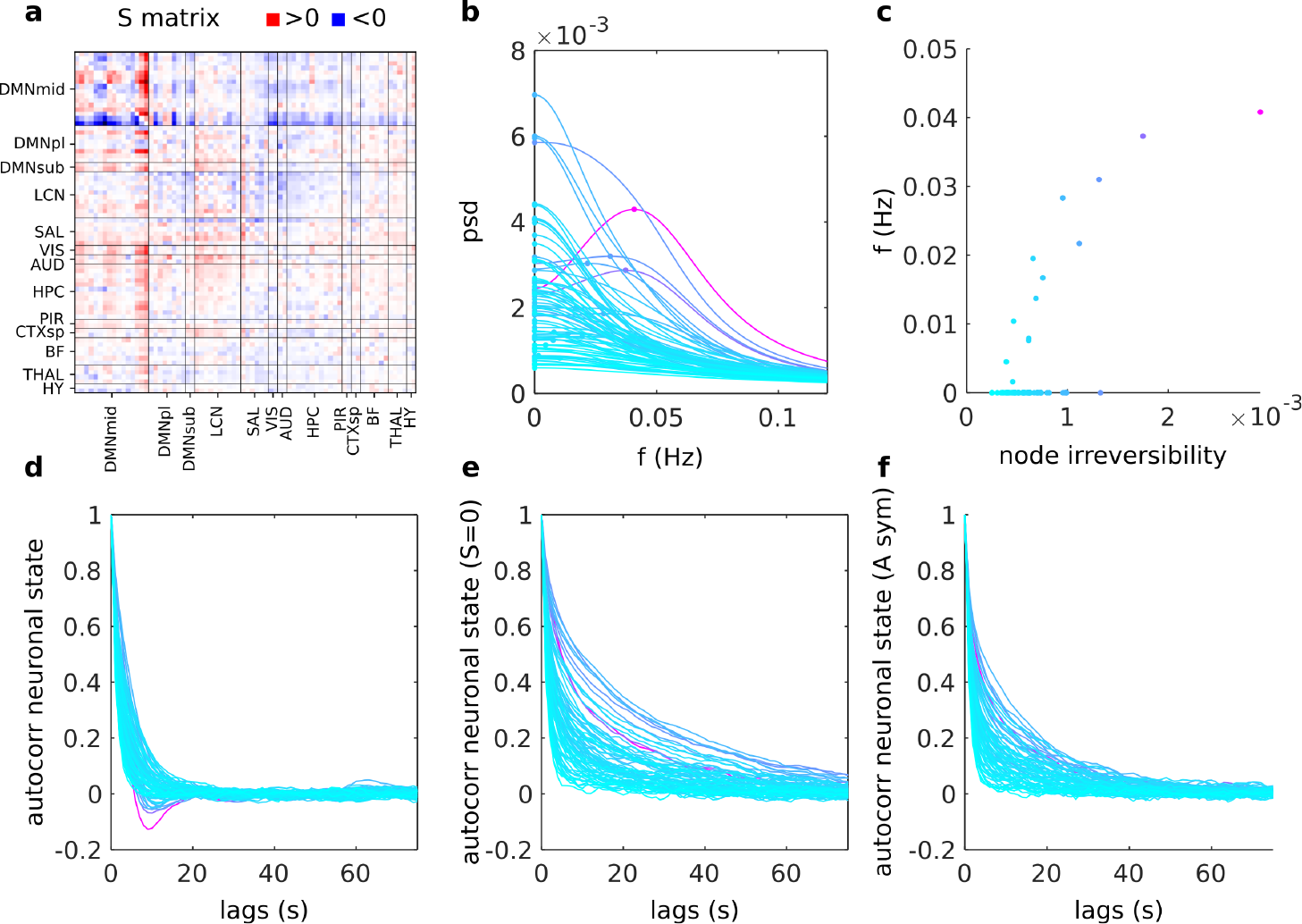
(a) The dC-Cov (*S* matrix) for a representative subject from the mouse dataset is displayed. Nodes are categorized based on functional networks. Positive entries are depicted in red, while negative entries are represented in blue. Following the adopted convention, a positive entry indicates that the node in the column acts as the source, while the node in the row serves as the target; (b) the power spectral density (psd) of each neuronal state, color identifies the level of irreversibility of the node from 0 (cyan) to higher values (purple); (c) for each node, how its level of irreversibility relates with the frequency of the psd peak; (d) the auto-correlation of each neuronal state; (e) the auto-correlation of each neuronal state computed by imposing *S* = 0 and deriving the corresponding symmetric *A* matrix; (f) the auto-correlation of each neuronal state inferred from a sparse-DCM model that was constrained to be time-reversable.

To gain a better grasp of how the dC-Cov influences steady-state dynamics, we examined the DCM model under three different situations. These include using the original data and the time-reversed data with the standard sparse-DCM model, and using the original data with a sparse-DCM designed to maintain symmetry in the *A* matrix.

For the mouse dataset, we initially presented two metrics that quantify the model’s capacity to replicate two key data features (static functional connectivity - sFC and dynamic functional connectivity - dFC, see Section 2.5.3) (see Fig. 2a) under the three previously mentioned conditions. This aims to assess the quality of the single-subject model inference by evaluating its capability of generating synthetic fMRI recordings that resemble the sFC and dFC of the empirical one. We observed no substantial distinctions between the inferences made on the original and time-reversed data. It is worth noting that this outcome was anticipated, as the metrics we employed are not sensitive to time-directionality. Moreover, it is important to mention that the model’s performance declined when symmetry was enforced in the *A* matrix.

**Figure 2.**
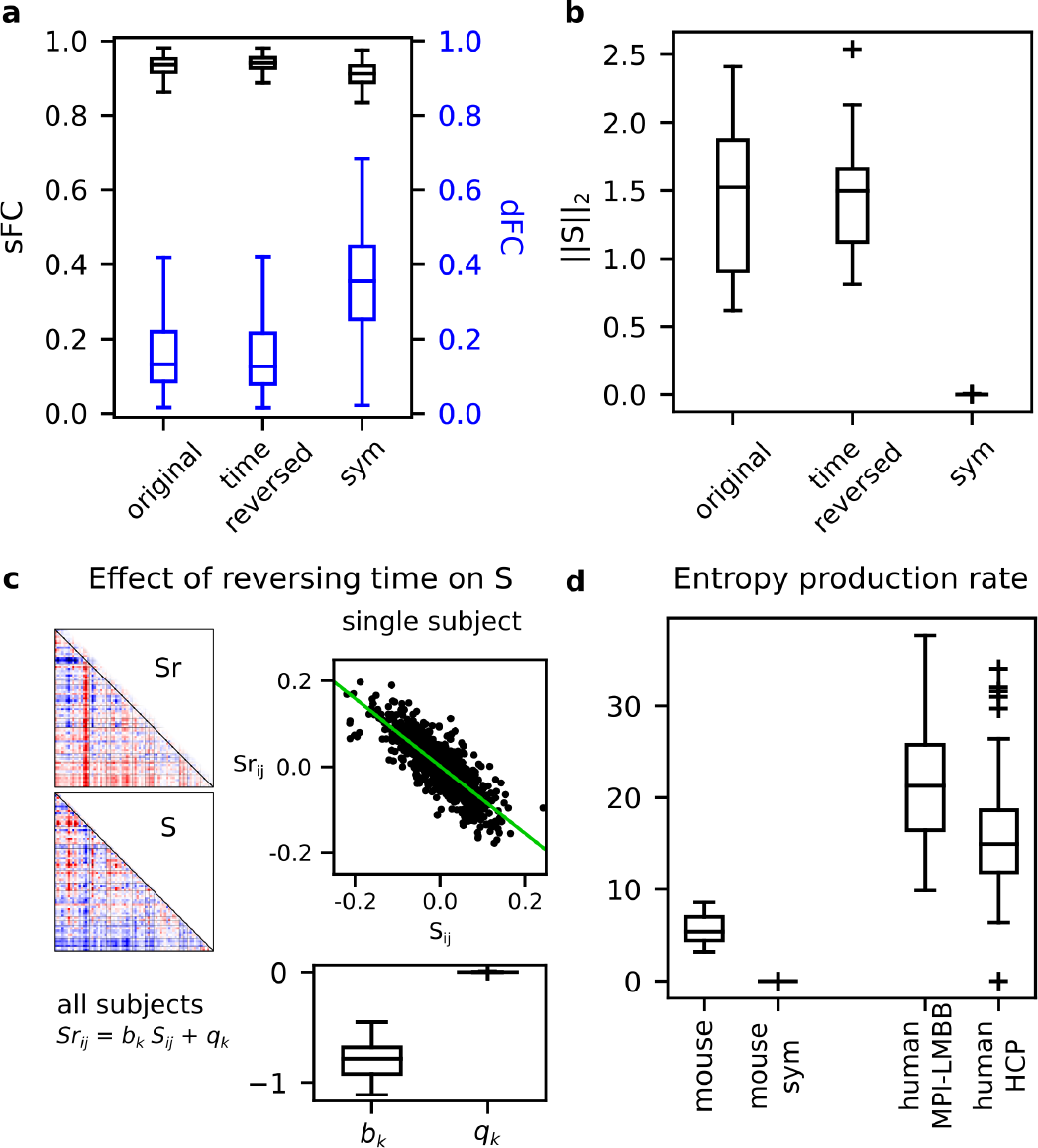
(a) The model’s ability to generate data akin to the empirical mouse recordings is shown in terms of the correlation between their static functional connectivity (sFC) (left y-axis) and the Kolmogorov-Smirnov (KS) distance of their dynamic functional connectivity (dFC) (blue right y-axis). This is demonstrated for the original data, time-reversed data, and the original data fitted with a DCM model designed to maintain symmetry in the effective connectivity (EC); (b) the 2-norm of the *S* matrix is depicted under the three inference conditions: original, time-reversed, and sym (symmetry); (c) the impact of reversing time on the dC-Cov is illustrated. For a representative subject, the scatter plot displays the expected negative correlation between corresponding entries of *S* and *S*_*r*_ (the *S* matrix inferred with time reversal). Additionally, the boxplots show the associated distributions of slope (*b*_*k*_) and intercept (*q*_*k*_) of the single-subject linear fit across the mouse dataset; (d) the entropy production rate computed in the mouse and human datasets.

In Fig. 2b, we reported the 2-norm of the *S* matrix under the three inference conditions. The *S* matrices inferred from the original and time-reversed data demonstrated similar 2-norm values, whereas the *S* matrix reduced to 0 when symmetry was enforced in the model. These outcomes align with theoretical expectations, which are also reflected in Fig. 2c, depicting the impact on the dC-Cov when reversing the time direction of recordings. Each entry of *S* should correspond to its opposite value in *S*_*r*_ (the *S* matrix inferred with time reversal). In panel c, the scatter plot illustrates the entry-wise relationship between *S* and *S*_*r*_ for a representative subject, while the boxplots display the distributions of slope (*b*_*k*_) and intercept (*q*_*k*_) across the mouse dataset. Finally, in panel (d), we compare the entropy production rates computed as described in Eq. 10 for each subject across datasets. Interestingly, both human datasets exhibited higher entropy production rates than the mouse dataset.

After establishing the reliability of our model’s inference in reproducing key FC features and reacting to signal time-reversion, we shifted our focus to examining changes in phase coherence among regions in the simulated data. The presence of alternating periods of co-activation and deactivation among regions is a fundamental characteristic of resting-state fMRI data. These dynamics can be quantified using synchronization and metastability indices, as described in Section 2.5.3. These measures stem from the Kuramoto order parameter and were initially developed to investigate phase transitions in models of coupled oscillators [71, 20]. Given the linear nature of our state model, the phase space possesses only a fixed point, limiting our ability to replicate complex dynamic phenomena. Despite this limitation, we found it valuable to explore eventual disparities between modeled and empirical data in terms of synchronization and metastability. We conducted this analysis using both the original model and its time-reversible version. Moreover, we examined the influence of the hemodynamic function on these measures. It is worth noting that across datasets, data generated from the subject-level sparse-DCM model closely mirrors the synchrony index of the empirical data. This alignment is evident in the top panels of Fig. 3, where the orange boxplot in the “state+hrf” columns closely corresponds to the gray boxplot representing the empirical BOLD signal. Likewise, the synchronization index of the time-reversible sparse-DCM (green boxplots) does not exhibit significant differences from the empirical data, indicating that asymmetry may not be necessary to replicate empirical synchronization. The scenario differs when it comes to metastability, where our simulations revealed a significant dependence on the dataset. In the case of mouse recordings, both time-reversible and time-irreversible models managed to replicate metastability, albeit with the time-reversible model (green boxplot) yielding a lower average metastability index. Conversely, for the MPI-LMBB dataset, only the time-irreversible model effectively reproduced empirical metastability, while neither model could accurately replicate metastability in the HCP dataset. In all cases, the time-reversible models deviated further from the empirical values. Finally, this analysis sheds light on the role of the hemodynamic response function (HRF) in modeling the BOLD signal. In particular, it highlights its capacity to influence both synchrony and metastability. Indeed, both indices increased if computed on data when the HRF was included.

**Figure 3.**
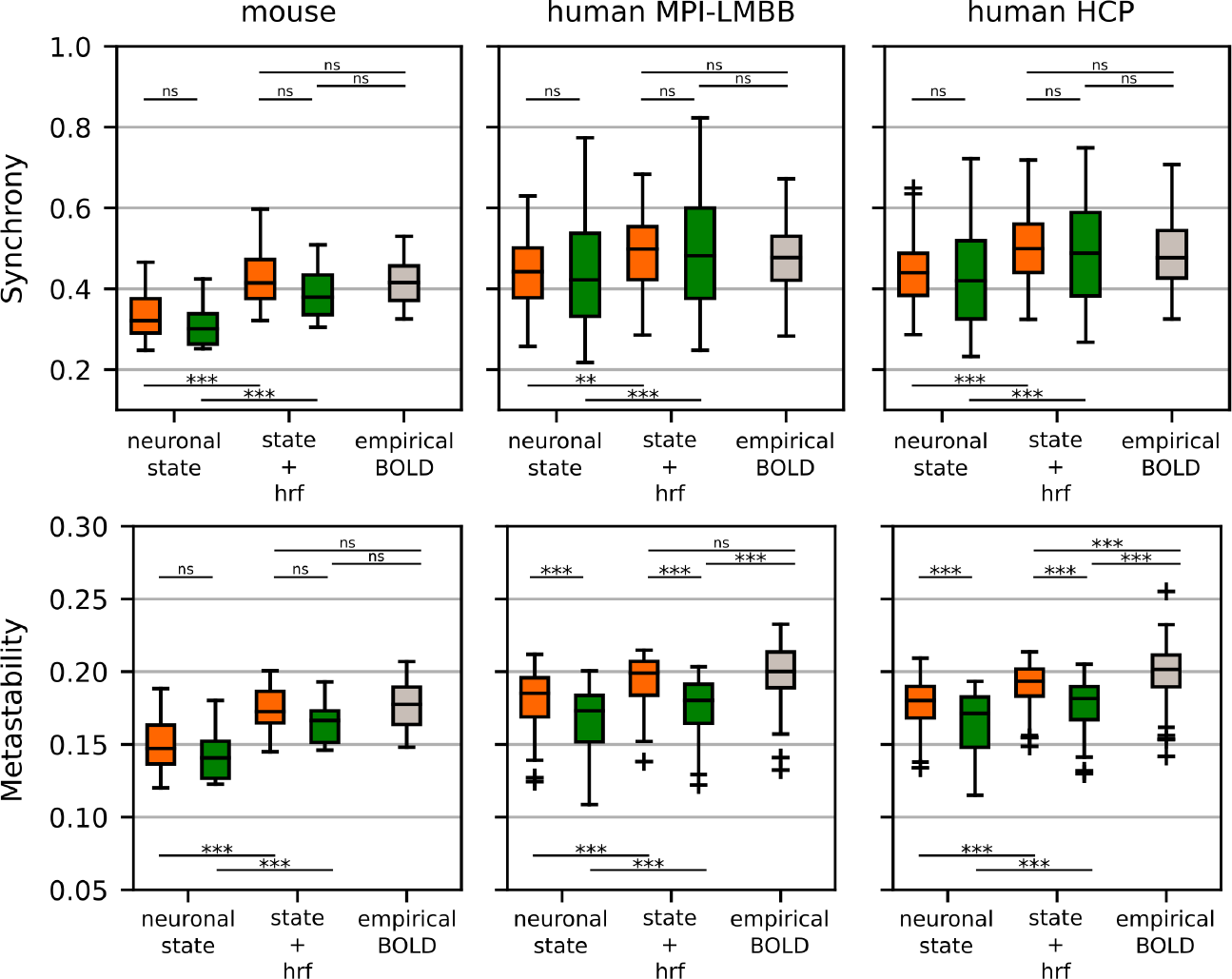
Synchrony and metastability indices in each dataset (mouse, human MPI-LMBB, and human HCP) using both simulated and empirical BOLD signals. When simulating the data, two factors were taken into account: the inclusion of the hemodynamic response function (HRF), labeled in the figure as “neuronal state” vs. “state+hrf”, and the time-reversibility of the model, i.e. time-irreversible (orange boxplots) vs. time-reversible (green boxplots). *\** = *p <* 0.05, *\*\** = *p <* 0.01, *\*\*\** = *p <* 0.001 and ns = not significant, ANOVA test with Tukey’s multiple comparison test.

The second part of the Results section focuses on how a non-zeros dC-Cov defines the directionality on which interactions across regions occur. The research was motivated by a question regarding the interpretability of the strengths of rows and columns in the EC matrix. Specifically, it aimed to determine whether it is valid to interpret the sums of rows and columns in the EC matrix as representing the incoming and outgoing connection strengths, respectively. In this context, our starting point was Eq. 4, with a particular focus on the statistical interpretation that each matrix obtained by decomposing *A* has in term of expectation. To recap, the diagonal matrix Σ_*w*_ contains the variance of the node random fluctuations (assumed constant across nodes), Σ represents the steady-state covariance matrix, denoted as 𝔼 [*x*(*t*)*x*(*t*)^*T*^] |_*t→∞*_, and its inverse Σ^*−*1^ is the precision matrix. This latter holds a direct interpretation in terms of conditional independence: a value of zero or very close to zero indicates that the two signals are conditionally independent given the rest of the recordings [19]. Finally, the steady-state dC-Cov *S* with off-diagonal entries 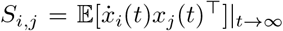 and zero diagonal, which facilitates the identification of source and sink nodes in each pair.

Beginning with the EC row strengths, it has been observed from Eq. 7 that the sum of its *i*-th row can be decomposed as a linear combination. This combination involves regressors that are equal to the row sums of the precision matrix Σ^*−*1^ and coefficients that are the elements in the *i*-th row of *S*. Additionally, there is the term 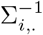, weighted by *−σ*^2^*/*2, which accounts for the dissipative contribution that is mediated by the dA-Cov.

In Fig. 4, the first row illustrates how *A* row sums relate with Σ^*−*1^ row sums (panel a) and with *S* row sums (panel b). In the case of the former, a strong negative correlation is evident. However, in the latter, there is no clear pattern. This suggests that in our resting-state BOLD recordings, even in the presence of a non-zero *S* matrix, the relationship between row sums of *A* and Σ^*−*1^ is primarily preserved. It is important to note that in the case of a time-reversible model, there would be a perfect linear relationship with a slope equal to *−σ*^2^*/*2, as indicated by the blue line in panel (a). While the influence of the dC-Cov does not emerge from the row sums of *A*.

**Figure 4.**
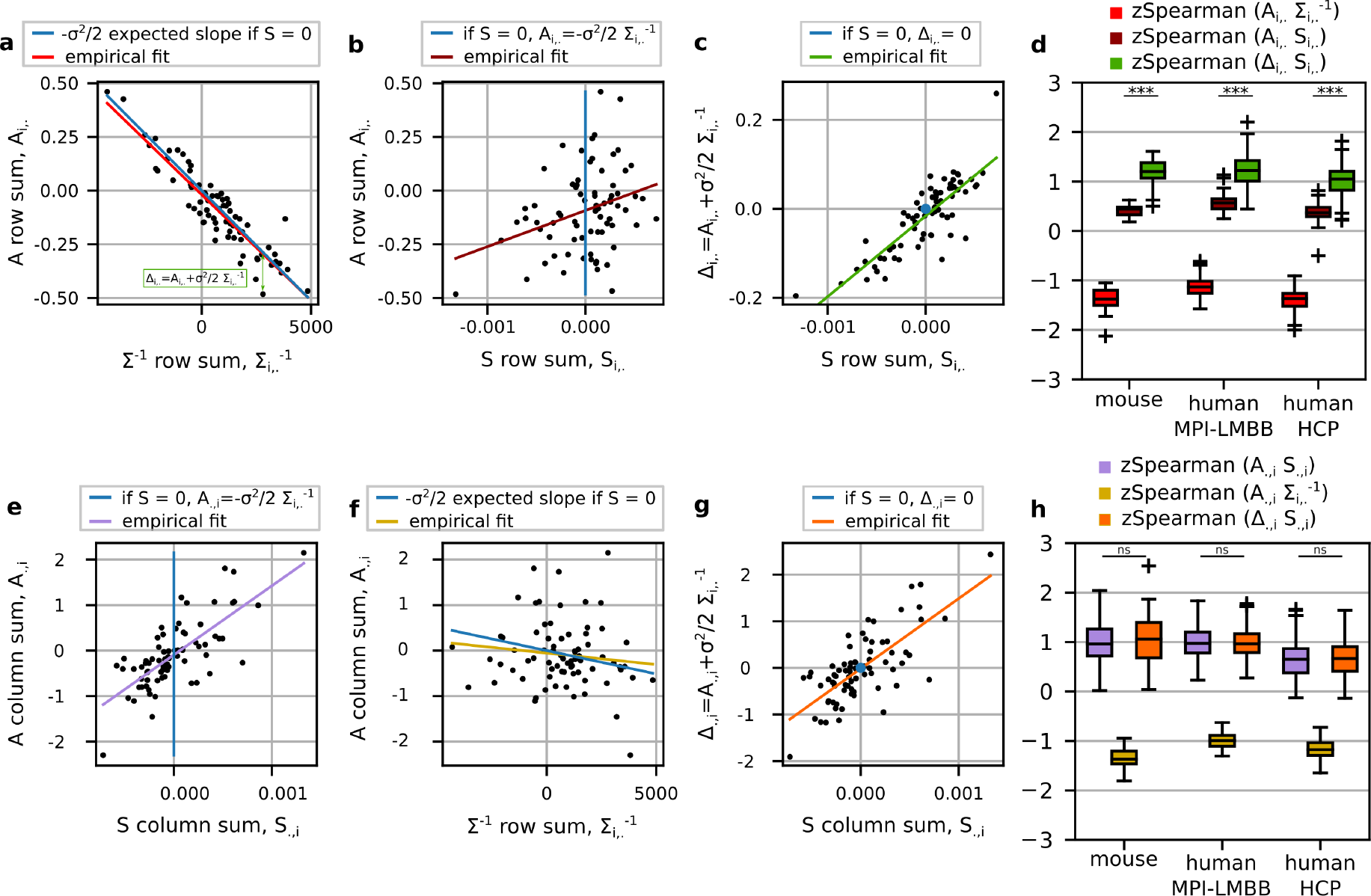
How the row and column strengths of the EC matrix (*A*) relate with the corresponding strengths computed on the matrices on which *A* can be decomposed, i.e. Σ^*−*1^ and *S*. Panels on the top refer to the EC row strengths: in panel (a) the relationship between the row sums of *A* and the corresponding row sum of Σ^*−*1^ (the regressors in Eq. 7), in panel (b) how *A* row strengths relate with the row sums of *S* (the coefficients in Eq. 7), in panel (c) *S* row strengths are shown in relation with Δ_*i*,._ which isolate the contribution of *S* in the row strength of *A*, and panel (d) summaries the previous relationships by showing the distributions of the Fisher transformed Spearman correlations across subjects for each dataset. Statistical t-test was performed to compare the use of *A* or Δ to reveal what encoded in the row sum of *S*. Δ performed significantly better, *p <* 0.001. Panels on the bottom refer to the EC column strengths: in panel (e) the relationship between the column sums of *A* and the corresponding column sums of *S* (the regressors in Eq. 8), in panel (f) how *A* column strengths relate with the node strengths of Σ^*−*1^ (the coefficients in Eq. 8), and panel (g) the column sum of Δ against the one of *S*. Panel (h) generalises the previous results across subjects and datasets, similarly to panel (d). No significant differences between using *A* or Δ.

To emphasize the effect of the dC-Cov, we need to isolate its contribution to the row sums of *A*, which shows how much each row sum deviates from the time-reversible condition. In Fig. 4(a), this is represented in green as 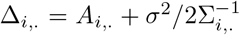, and is shown in a scatter plot against the row sum of *S* in panel (c). Now, the contribution of the solenoidal flow becomes apparent as these two variables linearly correlate. Note that Δ = *S*Σ^*−*1^ represents the gradient of the solenoidal flow.

The scatter plots in Fig. 4 pertain to a representative mouse subject. Comprehensive results across datasets are depicted in panels (d) and (h).

Regarding the EC column strengths, as reported in Eq. 8 for each node *i* its column sum is a linear combination with regressors the row sums of *S* and coefficients the *i*-th column of Σ^*−*1^, plus the term in Eq. 6. Fig. 4(e) illustrates the relationship between the column sums of *A* and the corresponding column sums of *S*. It appears that the column sums of *A* positively correlate with the corresponding column sums of *S*, indicating that the solenoidal part is reflected in the columns of *A*. In panel (f), we also explored the relationship between the column sums of *A* and the corresponding row sums of Σ^*−*1^. The expected relationship for a time-reversible model is represented by the blue line, but the empirical values do not reveal any significant trend between them. For the sake of completeness, in panel (g), we examined the relationship between the column sums of Δ and the column sums of *S*. Global findings presented in panel (h) validate that the influence of the dC-Cov at the node level can be inferred from the columns of the EC. This observation holds true whether or not the dissipative component is considered, as there were no significant differences between using *A* or Δ.

We conclude the Results section by showing the column sums of *S* in each dataset, the in/out node profile, and a reduced version of dC-Cov in which entries referring to the same pair of networks have been averaged (in/out network matrix), see Fig. 5. Considering the interpretation of the dC-Cov, its column strengths quantify the prevalence of a node acting as a source (positive strength) or a sink (negative strength) within the system. In panel (a), we present the in/out node profile and the in/out network matrix derived from the mouse dataset. Similarly in panels (b) and (c) for the MPI-LMBB and HCP datasets, respectively. The correlation coefficient between the two human profiles is 0.73 (*p <* 0.001) and between the in/out network matrices is 0.89 (*p <* 0.001), suggesting good reproducibility across datasets and enhancing the reliability of our results. In [70] a similar analysis was carried out using structural connectivity (SC) data. In the case of human data, they utilized undirected SC based on diffusion MRI, and for the mouse case, directed SC obtained from the connectome in [57]. The study focused on employing asymmetric network communication measures with the objective of identifying sender and receiver nodes, with which we compared our findings. In particular, the strongest correlations with our results were obtained when comparing with the asymmetric measure based on diffusion efficiency. The in/out network matrices in Fig. 5 exhibited Pearson correlation coefficients of 0.55 (*p* = 0.03) for the mouse dataset, 0.57 (*p <* 0.01) for the MPI-LMBB dataset, and 0.52 (*p* = 0.02) for the HCP dataset. Supplementary Figure S1 displays the coupling at the single-subject level with the diffusion efficiency asymmetry, across different SC densities, for each functional dataset. The couplings with diffusion efficiency were computed using both EC *−* EC^T^ and dC-Cov. A consistently stronger coupling with structural asymmetry was observed when using dC-Cov.

**Figure 5.**
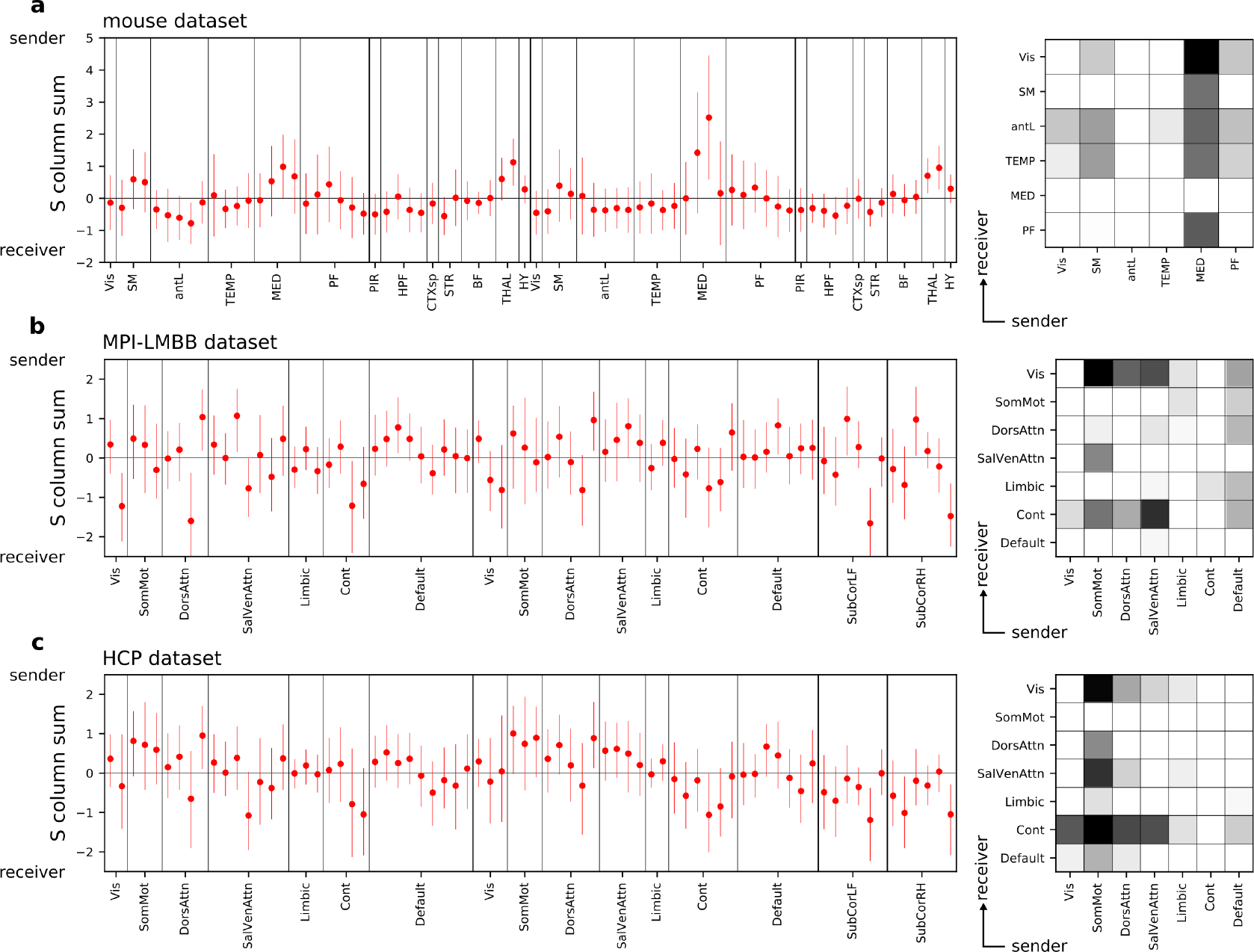
The average in/out node profile and in/out network matrix across subjects for each dataset are presented in the following panels. In the in/out network matrix, only positive values are displayed, with the grayscale colorbar transitioning from lighter to darker shades of gray to indicate that the network on the x-axis predominantly interacts as a sender with the network on the y-axis. In panel (a), the in/out node profile and network matrix are derived from the mouse data. Panels (b) and (c) display the in/out node profiles and network matrices of the MPI-LMBB and HCP datasets, respectively. For each node, a negative column sum in the matrix *S* indicates a prevalence of the node behaving as a receiver, while a positive column sum in *S* suggests a prevalence of the node behaving as a sender.

## 4 Discussion

In this work, we explored the implications of having a non-zero differential cross-covariance (dC-Cov) in the modeling of fMRI data using a linear state-space stochastic model. Specifically, we examine the DCM-fMRI framework, which incorporates an observational equation to model the hemodynamic transfer function, i.e. the forward model. DCM, initially employed for task-based recordings and more recently for resting-state data, is a well-established family of top-down models. In this framework, the state interaction matrix is referred to as the effective connectivity (EC) matrix. This matrix is known for its ability to convey directional and causal information through its asymmetric structure. This stands in contrast to functional connectivity (FC), which is symmetric. Consequently, EC has frequently been employed to differentiate between incoming and outgoing node connections. Indeed, while the asymmetry of EC is not a novel concept, there has been a relative lack of attention directed towards elucidating the underlying reasons for its asymmetric nature and the mechanistic implications when generating surrogate data utilizing a state matrix of such a form.

The asymmetry of EC is a consequence of a non-zero dC-Cov, and this is intricately linked with the presence of non-flat brain hierarchies. Therefore, the central question of this work focuses on how we can discern a non-flat brain hierarchy from the data, identify the patterns of brain regions involved, and ascertain the direction in which the hierarchy functions.

We showed that the crucial element for addressing these questions is the steady-state dC-Cov. We derived it at the single-subject level by identifying a stochastic DCM model using an inference method known as sparse-DCM [62]. In particular, our study utilized a matrix decomposition that enables the separation of the zero-lag statistic from the non-zero-lag statistic, and highlighted the role of the latter component which imparts the asymmetric shape to the state matrix. Interestingly, when considering a stochastic model near a stable fixed point with Gaussian fluctuations [48], this decomposition can be interpreted from the perspective of statistical physics as representing the splitting of the non-quilibrium steady-state landscape into two orthogonal components: the gradient of the potential energy (a dissipative gradient flow) and the curl steady-state flow (a solenoidal flow) [27].

The curl steady-state flow which in our modeling framework is mediated by the dC-Cov, is a manifestation of a non-equilibrium dynamics. Non-equilibrium dynamics have been found to be ubiquitous among living systems [35], and they also manifest within rsfMRI recordings [46].

A non-zero dC-Cov has several effects on the data. Firstly, it underscores the significance of time-delayed interactions, which have been consistently highlighted as playing a central role in rsfMRI data. For example, in [72], it has been shown how both time and spatial autocorrelation can account for a significant portion of the network topological measures commonly used in connectome analyses; in [11], the significance of delayed interactions has been highlighted when modeling neural data; additionally, in [5], the connection between various empirical phenomena observed in large-scale brain organization and both zero-lag and time-lag statistics has been explored. Moreover, time-delayed interactions define the so-called arrow-of-time [6, 22], which is inherently linked to the concept of entropy production rate, the center of recent works, e.g. [52, 47].

Interestingly, as previously reported in the literature, a time-irreversible dynamics can give rise to slow fluctuations due to the presence of oscillatory modes represented by the complex eigenvalues of the Jacobian matrix, see Fig. 1. It is reasonable to assume that these oscillatory features, spanning various timescales and spatial patterns, are linked to or can be considered as linear projections (in the case of non-linearity) of phenomena such as harmonic waves [2], traveling waves [5], oscillatory modes [10], brain spirals [80], and cascade propagation [63].

Examining the contribution of each brain region to the definition of the curl steady-state flow reveals its role in shaping how information propagates throughout the brain hierarchy and its overall tendency to primarily function as a sender or a receiver. The notion of brain hierarchy is widely recognized in systems neuroscience, as highlighted in [42], Of interest for this discussion is the functional cortical hierarchy, which reflects graph structural properties of the functional connectivity matrix [54]. In recent years, significant progress has been made in understanding functional connectomics, including the identification of major functional gradients that explain key aspects of functional connectivity. However, there has been a gap in understanding the directional aspects of information propagation along these gradients.

The modeling approach we have employed in this study simultaneously addresses the need to consider both the zero-lag statistic, which represents functional connectivity, and the non-zero lag statistic, which corresponds to differential covariance. The diagonal entries of the differential covariance contains information regarding how each node influences its own derivative (dA-Cov) while the off-diagonal part represents the causal component that describes how and to what extent each brain region influences the temporal dynamics (i.e., the derivative) of others. All these elements are integrated within the effective connectivity matrix, which has the capacity to define all trajectories within the state space that adhere to the previously mentioned statistics.

This highlights the pivotal role of the model identification algorithm, as its performance significantly influences the ability to accurately reproduce empirical data statistics. In our study, we employed a recently developed approach to address the inference problem of a DCM model for rsfMRI. This approach incorporates two critical components: a sparsity-inducing prior applied to the effective connectivity matrix and a linearized region-specific hemodynamic response function (HRF) [62].

Part of our results, particularly those presented in Figs. 2 and 3, were intended to provide evidence regarding the model identification process. We focused on standard metrics commonly used in the literature for this purpose. They include the model’s ability to replicate the empirical static FC and the range over which dynamic FC values are distributed. These metrics were computed following the inference of three different models for each subject. The first model applied the standard sparse-DCM to the original data. The second model applied the standard sparse-DCM to the time-reversed data, aiming to illustrate how this impacts the steady-state dC-Cov. The third model utilized a time-reversible sparse-DCM. Collectively, this comparison demonstrated that the inferred models generally exhibited good generalization properties, except for the time-reversible model. This finding supports the idea that a non-equilibrium dynamics model provides a better explanation for rsfMRI data.

We also evaluated other metrics, specifically the synchrony and metastability indices. Although these metrics may seem at odds with the stable single fixed point around which the dynamics were modeled, as they are associated with nonlinear phenomena, we deemed it important to assess the disparities between surrogate and empirical data. Notably, only the metastability of the HCP dataset was not replicated by the surrogate data, while in all other cases, the simulated data exhibited indices that were not significantly different from the empirical ones.

We should examine this evidence from two perspectives. On one side, it is important to consider the specific characteristics of each dataset, such as the use of halothane anesthetic in the mouse data, variations in sampling rates across datasets, and notably, the fact that the HCP dataset has the lowest sampling rate. On the other side, we should also acknowledge that recent research has reported the ability of a linear state-space model to replicate large-scale brain dynamics successfully [56, 74, 61].

In this context, it is worth mentioning the advantages offered by the linear state-space model when computing the steady-state dC-Cov and metrics related to time-irreversibility, such as the entropy production rate, for which closed-form solutions are available [32]. Alternative approaches have been proposed and tested for estimating dC-Cov in previous researches, e.g. [49, 15, 45]. Additionally, in [52], a hierarchical clustering algorithm was used to sample the system phase space and derive the entropy production rate, while [21] introduced an estimator of time-irreversibility by comparing the correlation between forward and backward signal lag-correlations.

The second part of the Results section delved into the issue of interpretation. Specifically, we explored whether the EC matrix or the steady-state dC-Cov offers a more appropriate description of how information flows within the brain hierarchy.

Traditionally, the EC matrix has been used to reveal directed connections within the brain [25]. However, when we consider the role of EC as a state-space matrix, it provides timing information by modeling the trajectory of the state space for each initial condition. In doing so this matrix combines various data statistics, including the variance of noise fluctuations, the zero-lag covariance, and the dC-Cov, as seen in Eq. 4. On the other hand, the steady-state dC-Cov serves a distinct purpose. Firstly, it isolates the causal component of the EC (what makes EC asymmetric). Additionally, it acts as rotation matrix so it mediates the solenoidal component of the landscape (what contributes to making the hierarchy non-flat). Note that it is not enough to separate the asymmetric part of EC since this could be done also by computing EC *−* EC^T^ [29], however this separation of the asymmetric part does not represent the steady-state dC-Cov.

In accordance with the conventional definition of EC incoming node strength, our empirical observations highlighted the substantial impact of the precision matrix in estimating a node’s incoming strength. The influence of the dC-Cov, as manifested in the effective connectivity matrix, became evident primarily after the dominant contribution of the precision matrix had been factored out. This relationship is elucidated in the first row of Fig. 4. Conversely, when calculating the outgoing node strength from EC, a consistent linear relationship with the dC-Cov was maintained, as illustrated in Fig. 4‘s second row. Thus suggesting for this latter a coherent interpretation in terms of directionality. Building upon the interpretation of the dC-Cov, we devised a straightforward metric to encapsulate each node’s overall role within the network. This metric involved summing the columns of the dC-Cov matrix for each node, obtaining what we called in the Result section the average in/out node profile. A positive sum indicated that the node primarily acted as a source, whereas a negative sum suggested a sink role. It is important to note that by computing the row sums and reversing the interpretation, we would arrive at the same conclusions. Likewise, instead of reducing the role of each node to a single number, which would overlook network heterogeneity, we also presented the in/out network matrix. In this matrix, entries of the dC-Cov matrix were averaged, providing insight into how each pair of networks interacts in terms of source or sink behavior.

Concerning the mouse dataset, a predominant sender role was assigned to the medial (MED), prefrontal (PF) and somatomotor (SM) regions which constitute the DMNmid and LCN functional networks. This aligns with findings from [17] where these regions exhibited a high out/in structural connection ratio. While a receiver role at the cortex level is mainly played by anterolater (antL) areas of which its main component is the agranular insular area which forms the salience network. Additionally, our results revealed the thalamic and hypothalamic regions as prominent senders, even though a specific preferred directionality was not previously reported in the cited work but however identified as global connector hubs. As main subcortical receivers, we identified the hippocampus, and the dorsal striatum (DMNsub network) which was reported as main receiver also in [17] being part of the basal ganglia.

In relation to the human datasets, we firstly highlight the high degree of consistency between the corresponding results. This consistency may be attributed in part to the use of the same parcellation scheme. Nevertheless, it adds robustness to our findings, as it indicates that similar results were obtained from two separate independent datasets. Additionally, our results align with [70], which also explored the asymmetric behavior of brain regions in terms of being a sink or source using network communication measures on structural imaging data (diffusion MRI). Specifically, our results support the concept that information travels through the functional hierarchy, progressing from sensory/motor and unimodal regions to integrative multimodal ones responsible for higher-order functions. This explains the dominant sender activity of the somatomotor network, primarily communicating with the visual, ventral attention, and control networks. And similarly, the connections from the ventral attention network to the visual and control networks, as well as the role of the control network as a primary receiver from the other networks. The widespread agreement in the literature regarding this directional arrow is substantiated by evidence from various sources. These include studies on structural data [55, 70], investigations into functional gradients [54], and dynamic models [24, 23, 58]. According to this line of works, the default mode network should occupy the apex of the hierarchy and primarily receive inputs from all other networks. However, our findings reveal a different pattern. In the MPI-LMBB dataset, the default mode network takes on a sender role, while in the HCP dataset, it does not seem to exhibit a distinct preferred behavior. This observation is in concordance with what hypothesised by the predictive processing framework. This framework postulates the existence of two counteracting information flows within the brain: top-down prediction signals (feedback signals) and bottom-up prediction errors (feedforward signals) [13, 44]. It is plausible that the dominant behavior of the default mode network in our results can be attributed to the prevalence of top-down prediction signals originating from it.

A similar analysis based on the asymmetry of EC was conducted also in [70]. In their analysis, asymmetry was computed by subtracting EC from its transpose. However, as previously mentioned, this approach captures the asymmetric part of EC but does not represent the dC-Cov. In their comparison using a cortical partition into 7 networks (the same partition used in our analysis), they did not observe a significant correlation between the asymmetry derived from EC and the asymmetry derived from communication-based measures. Similarly, our dC-Cov showed no significant correlation with the EC asymmetry reported in [70], whereas it did correlate with the asymmetry derived from diffusion MRI data.

Future research will delve deeper into the exploration of the steady-state dC-Cov within the framework we have presented here. One crucial aspect, which we have yet to explore in this study but believe holds significant importance, is the differentiation of dC-Cov across various timescales. It might be overly simplistic to assume that the dynamics of large-scale brain activity are governed by a single hierarchy. Evidence in favor of this notion can be found in [76], which elucidates the interaction between top-down and bottom-up processes. Additionally, in [8] authors demonstrated the intricate interaction between gradient-specific dynamics, and dynamics across gradients. And, in [14], researchers leveraged eigen-microstate analysis to identify basic modes in rsfMRI data. However, in most of these examples, a low-dimensional representation of brain activity was found, and each dimension characterized by its specific pattern of interactions but what is missing is the consideration of directionality. This could be studied as continuation of this work by applying an eigen-mode decomposition to the effective connectivity matrix and projecting the dC-Cov onto each mode. This could help disentangle the brain hierarchy across different timescales.

We conclude by emphasizing the importance of working at the single-subject level, a capability offered by the framework we utilized, although we primarily presented population-level results. This is a point that we will carefully consider in future work. Recent studies have demonstrated the relationship between the level of asymmetry in brain hierarchy and brain states evoked by task conditions [52], levels of conscious awareness [60], neuropsychiatric diseases [22], and the presence of stroke lesions [43]. This suggests the potentiality of deriving biomarkers based on properties of brain hierarchies.

## 5 Acknowledgements

This research was supported by the DEI Proactive grant “Personalized whole brain models for neuroscience: inference and validation” from the Department of Information Engineering of the University of Padova (Italy).

Data were provided in part by the Human Connectome Project, WU-Minn Consortium (Principal Investigators: David Van Essen and Kamil Ugurbil; 1U54MH091657) funded by the 16 NIH Institutes and Centers that support the NIH Blueprint for Neuroscience Research; and by the McDonnell Center for Systems Neuroscience at Washington University.

We thank A. Gozzi and L. Coletta from the Functional Neuroimaging Laboratory (Center for Neuroscience and Cognitive Systems @ UniTn, Istituto Italiano di Tecnologia, Rovereto, Italy) for providing the mouse dataset and offering assistance in data curation and preprocessing.

## Supplementary Material

**Table S1.**
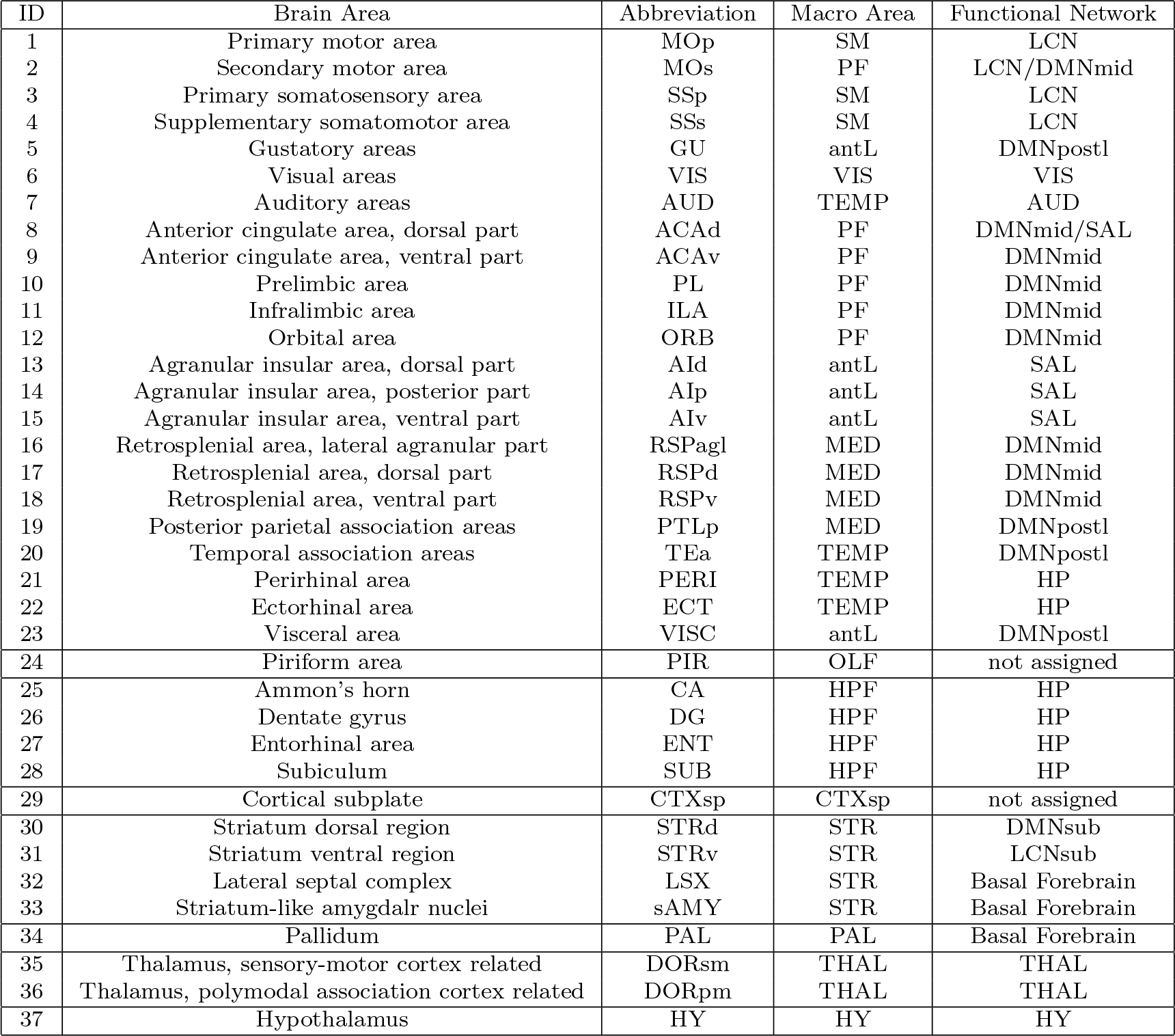
Mouse brain areas and network information. The same parcellation was applied on both hemispheres. Abbreviations used in the Macro Area column refer to the following areas: somatomotor (SM), medial (MED), temporal (TEMP), visual (VIS), anterolateral (antL), prefrontal (PF) and hippocampal formation (HPF). While abbreviations in the Functional Network column refer to: laterocortical (LCN), subcortical laterocortical (LCNsub), default mode midline (DMNmid), default model posterolateral (DMPpostl), default model subcortical (DMNsub), salience (SAL) and hippocampus (HP).

**Table S2.**
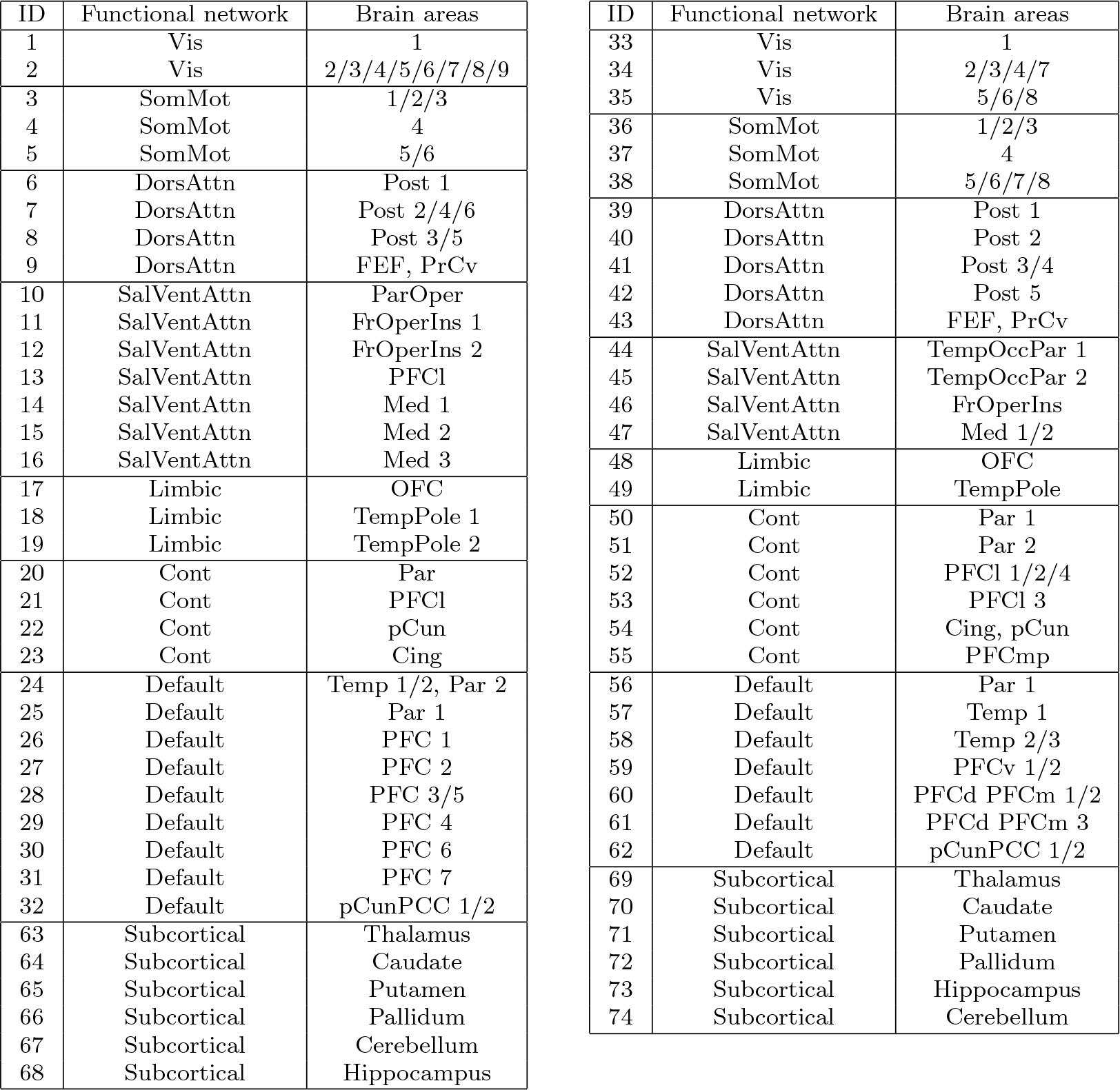
Human brain parcellation and network information (left table, left hemisphere; right table, right hemisphere) This is the result of a consensus clustering algorithm applied to the 100-area cortex parcellation (7-networks) of [68] and the AAL2 subcortical parcellation of [66].

**Figure S1.**
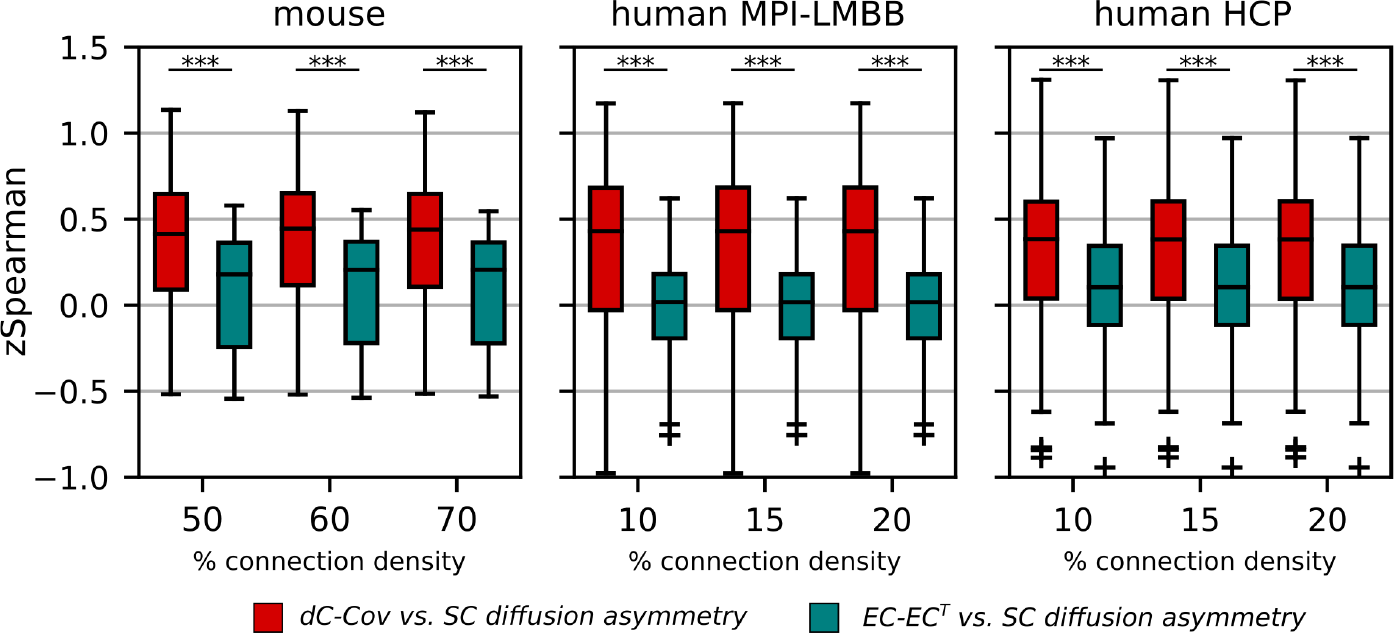
The coupling at the single-subject level with the diffusion efficiency asymmetry, across different SC densities, for each functional dataset. The couplings with diffusion efficiency were computed using both the dC-Cov (red) and EC *−* EC^T^ (green). A consistently stronger coupling with structural asymmetry was observed when using dC-Cov. Paired t-test, *p <* 0.001.

## References

[1] N. Asadi, I. R. Olson, and Z. Obradovic. The backbone network of dynamic functional connectivity. Network Neuroscience, 5:851–873, 11 2021.

[2] S. Atasoy, I. Donnelly, and J. Pearson. Human brain networks function in connectome-specific harmonic waves. Nature Communications 2016 7:1, 7:1–10, 1 2016.

[3] A. Babayan, M. Erbey, D. Kumral, J. D. Reinelt, A. M. Reiter, J. Röbbig, H. L. Schaare, M. Uhlig, A. Anwander, P. L. Bazin, A. Horstmann, L. Lampe, V. V. Nikulin, H. Okon-Singer, S. Preusser, A. Pampel, C. S. Rohr, J. Sacher, A. Thöne-Otto, S. Trapp, T. Nierhaus, D. Altmann, K. Arelin, M. Blöchl, E. Bongartz, P. Breig, E. Cesnaite, S. Chen, R. Cozatl, S. Czerwonatis, G. Dambrauskaite, M. Dreyer, J. Enders, M. Engelhardt, M. M. Fischer, N. Forschack, J. Golchert, L. Golz, C. A. Guran, S. Hedrich, N. Hentschel, D. I. Hoffmann, J. M. Huntenburg, R. Jost, A. Kosatschek, S. Kunzendorf, H. Lammers, M. E. Lauckner, K. Mahjoory, A. S. Kanaan, N. Mendes, R. Menger, E. Morino, K. Näthe, J. Neubauer, H. Noyan, S. Oligschläger, P. Panczyszyn-Trzewik, D. Poehlchen, N. Putzke, S. Roski, M. C. Schaller, A. Schieferbein, B. Schlaak, R. Schmidt, K. J. Gorgolewski, H. M. Schmidt, A. Schrimpf, S. Stasch, M. Voss, A. Wiedemann, D. S. Margulies, M. Gaebler, and A. Villringer. A mind-brain-body dataset of mri, eeg, cognition, emotion, and peripheral physiology in young and old adults. Scientific Data 2019 6:1, 6:1–21, 2 2019.

[4] G. Baron, E. Silvestri, D. Benozzo, A. Chiuso, and A. Bertoldo. Revealing the spatial pattern of brain hemodynamic sensitivity to healthy aging through sparse dcm. bioRxiv, page 2023.10.16.562585, 10 2023.

[5] T. Bolt, J. S. Nomi, D. Bzdok, J. A. Salas, C. Chang, B. T. T. Yeo, L. Q. Uddin, and S. D. Keilholz. A parsimonious description of global functional brain organization in three spatiotemporal patterns. Nature Neuroscience, 25:1093–1103, 8 2022.

[6] T. A. Bolton, D. V. D. Ville, E. Amico, M. G. Preti, and R. Liégeois. The arrow-of-time in neuroimaging time series identifies causal triggers of brain function. Human Brain Mapping, 44:4077–4087, 7 2023.

[7] M. Breakspear. Dynamic models of large-scale brain activity. Nature Neuroscience, 20:340–352, 3 2017.

[8] J. A. Brown, A. J. Lee, L. Pasquini, and W. W. Seeley. A dynamic gradient architecture generates brain activity states. NeuroImage, 261:119526, 11 2022.

[9] E. Bullmore and O. Sporns. Complex brain networks: graph theoretical analysis of structural and functional systems. Nature reviews. Neuroscience, 10:186–198, 3 2009.

[10] J. Cabral, F. Castaldo, J. Vohryzek, V. Litvak, C. Bick, R. Lambiotte, K. Friston, M. L. Kringelbach, and G. Deco. Metastable oscillatory modes emerge from synchronization in the brain spacetime connectome. Communications Physics, 5:184, 7 2022.

[11] F. Castaldo, F. P. dos Santos, R. C. Timms, J. Cabral, J. Vohryzek, G. Deco, M. Woolrich, K. Friston, P. Verschure, and V. Litvak. Multi-modal and multi-model interrogation of large-scale functional brain networks. NeuroImage, 277:120236, 8 2023.

[12] U. Casti, G. Baggio, D. Benozzo, S. Zampieri, A. Bertoldo, and A. Chiuso. Dynamic brain networks with prescribed functional connectivity, 2023.

[13] C. C. Chen, R. N. Henson, K. E. Stephan, J. M. Kilner, and K. J. Friston. Forward and backward connections in the brain: A dcm study of functional asymmetries. NeuroImage, 45:453–462, 4 2009.

[14] X. Chen, H. Ren, Z. Tang, K. Zhou, L. Zhou, Z. Zuo, X. Cui, X. Chen, Z. Liu, Y. He, and X. Liao. Leading basic modes of spontaneous activity drive individual functional connectivity organization in the resting human brain. Communications Biology, 6:892, 8 2023.

[15] Y. Chen, Q. Bukhari, T. W. Lin, and T. J. Sejnowski. Functional connectivity of fmri using differential covariance predicts structural connectivity and behavioral reaction times. Network Neuroscience, 6:614–633, 6 2022.

[16] L. Cocchi, L. L. Gollo, A. Zalesky, and M. Breakspear. Criticality in the brain: A synthesis of neurobiology, models and cognition. Progress in Neurobiology, 158:132–152, 11 2017.

[17] L. Coletta, M. Pagani, J. D. Whitesell, J. A. Harris, B. Bernhardt, and A. Gozzi. Network structure of the mouse brain connectome with voxel resolution. Science advances, 6, 12 2020.

[18] L. D. Costa, K. Friston, C. Heins, and G. A. Pavliotis. Bayesian mechanics for stationary processes. Proceedings of the Royal Society A: Mathematical, Physical and Engineering Sciences, 477, 12 2021.

[19] A. Das, A. L. Sampson, C. Lainscsek, L. Muller, W. Lin, J. C. Doyle, S. S. Cash, E. Halgren, and T. J. Sejnowski. Interpretation of the precision matrix and its application in estimating sparse brain connectivity during sleep spindles from human electrocorticography recordings. Neural Computation, 29:603–642, 3 2017.

[20] G. Deco and M. L. Kringelbach. Metastability and coherence: Extending the communication through coherence hypothesis using a whole-brain computational perspective. Trends in Neurosciences, 39:125–135, 3 2016.

[21] G. Deco, Y. S. Perl, H. Bocaccio, E. Tagliazucchi, and M. L. Kringelbach. The insideout framework provides precise signatures of the balance of intrinsic and extrinsic dynamics in brain states. Communications Biology 2022 5:1, 5:1–13, 6 2022.

[22] G. Deco, Y. S. Perl, L. de la Fuente, J. D. Sitt, B. T. T. Yeo, E. Tagliazucchi, and M. L. Kringelbach. The arrow of time of brain signals in cognition: Potential intriguing role of parts of the default mode network. Network Neuroscience, 7:966–998, 10 2023.

[23] G. Deco, D. Vidaurre, and M. L. Kringelbach. Revisiting the global workspace orchestrating the hierarchical organization of the human brain. Nature Human Behaviour 2021 5:4, 5:497–511, 1 2021.

[24] M. Demirtaş, J. B. Burt, M. Helmer, J. L. Ji, B. D. Adkinson, M. F. Glasser, D. C. V. Essen, S. N. Sotiropoulos, A. Anticevic, and J. D. Murray. Hierarchical heterogeneity across human cortex shapes large-scale neural dynamics. Neuron, 101:1181–1194.e13, 3 2019.

[25] L. Entz, E. Tóth, C. J. Keller, S. Bickel, D. M. Groppe, D. Fabó, L. R. Kozák, L. Eross, I. Ulbert, and A. D. Mehta. Evoked effective connectivity of the human neocortex. Human brain mapping, 35:5736–5753, 12 2014.

[26] D. C. V. Essen, K. Ugurbil, E. Auerbach, D. Barch, T. E. Behrens, R. Bucholz, A. Chang, L. Chen, M. Corbetta, S. W. Curtiss, S. D. Penna, D. Feinberg, M. F. Glasser, N. Harel, A. C. Heath, L. Larson-Prior, D. Marcus, G. Michalareas, S. Moeller, R. Oostenveld, S. E. Petersen, F. Prior, B. L. Schlaggar, S. M. Smith, A. Z. Snyder, J. Xu, and E. Yacoub. The human connectome project: a data acquisition perspective. NeuroImage, 62:2222–2231, 10 2012.

[27] K. J. Friston, E. D. Fagerholm, T. S. Zarghami, T. Parr, I. Hipólito, L. Magrou, and A. Razi. Parcels and particles: Markov blankets in the brain. Network Neuroscience, 5:211–251, 2 2021.

[28] K. J. Friston, L. Harrison, and W. Penny. Dynamic causal modelling. NeuroImage, 19:1273–1302, 8 2003.

[29] K. J. Friston, B. Li, J. Daunizeau, and K. E. Stephan. Network discovery with dcm. NeuroImage, 56:1202–1221, 6 2011.

[30] K. J. Friston, A. Mechelli, R. Turner, and C. J. Price. Nonlinear responses in fmri: The balloon model, volterra kernels, and other hemodynamics. NeuroImage, 12, 2000.

[31] S. Frässle, E. I. Lomakina, A. Razi, K. J. Friston, J. M. Buhmann, and K. E. Stephan. Regression dcm for fmri. NeuroImage, 3 2017.

[32] M. Gilson, E. Tagliazucchi, and R. Cofré. Entropy production of multivariate ornstein-uhlenbeck processes correlates with consciousness levels in the human brain. Physical Review E, 107:024121, 2 2023.

[33] M. Gilson, G. Zamora-López, V. Pallarés, M. H. Adhikari, M. Senden, A. T. Campo, D. Mantini, M. Corbetta, G. Deco, and A. Insabato. Model-based whole-brain effective connectivity to study distributed cognition in health and disease. Network Neuroscience, 4:338–373, 1 2020.

[34] M. F. Glasser, S. N. Sotiropoulos, J. A. Wilson, T. S. Coalson, B. Fischl, J. L. Andersson, J. Xu, S. Jbabdi, M. Webster, J. R. Polimeni, D. C. V. Essen, and M. Jenkinson. The minimal preprocessing pipelines for the human connectome project. NeuroImage, 80:105–124, 10 2013.

[35] F. S. Gnesotto, F. Mura, J. Gladrow, and C. P. Broedersz. Broken detailed balance and non-equilibrium dynamics in living systems: a review. Reports on Progress in Physics, 81:066601, 4 2018.

[36] J. Gonzalez-Castillo, I. S. Fernandez, K. C. Lam, D. A. Handwerker, F. Pereira, and P. A. Bandettini. Manifold learning for fmri time-varying functional connectivity. Frontiers in Human Neuroscience, 17, 7 2023.

[37] A. S. Greene, C. Horien, D. Barson, D. Scheinost, and R. T. Constable. Why is everyone talking about brain state? Trends in Neurosciences, 46:508–524, 7 2023.

[38] D. Gutierrez-Barragan, M. A. Basson, S. Panzeri, and A. Gozzi. Infraslow state fluctuations govern spontaneous fmri network dynamics. Current Biology, 29, 2019.

[39] D. Gutierrez-Barragan, N. A. Singh, F. G. Alvino, L. Coletta, F. Rocchi, E. D. Guzman, A. Galbusera, M. Uboldi, S. Panzeri, and A. Gozzi. Unique spatiotemporal fmri dynamics in the awake mouse brain. Current biology : CB, 32:631–644.e6, 2 2022.

[40] J. A. Harris, S. Mihalas, K. E. Hirokawa, J. D. Whitesell, H. Choi, A. Bernard, P. Bohn, S. Caldejon, L. Casal, A. Cho, A. Feiner, D. Feng, N. Gaudreault, C. R. Gerfen, N. Graddis, P. A. Groblewski, A. M. Henry, A. Ho, R. Howard, J. E. Knox, L. Kuan, X. Kuang, J. Lecoq, P. Lesnar, Y. Li, J. Luviano, S. McConoughey, M. T. Mortrud, M. Naeemi, L. Ng, S. W. Oh, B. Ouellette, E. Shen, S. A. Sorensen, W. Wakeman, Q. Wang, Y. Wang, A. Williford, J. W. Phillips, A. R. Jones, C. Koch, and H. Zeng. Hierarchical organization of cortical and thalamic connectivity. Nature 2019 575:7781, 575:195–202, 10 2019.

[41] S. Heitmann and M. Breakspear. Putting the “dynamic” back into dynamic functional connectivity. Network Neuroscience, 2:150–174, 6 2018.

[42] C. C. Hilgetag and A. Goulas. ‘hierarchy’ in the organization of brain networks. Philosophical Transactions of the Royal Society B: Biological Sciences, 375:20190319, 4 2020.

[43] S. Idesis, M. Allegra, J. Vohryzek, Y. S. Perl, J. Faskowitz, O. Sporns, M. Corbetta, and G. Deco. A low dimensional embedding of brain dynamics enhances diagnostic accuracy and behavioral prediction in stroke. Scientific Reports 2023 13:1, 13:1–17, 9 2023.

[44] Y. Katsumi, J. E. Theriault, K. S. Quigley, and L. F. Barrett. Allostasis as a core feature of hierarchical gradients in the human brain. Network Neuroscience, 6:1010–1031, 10 2022.

[45] T. Kim, D. Chen, P. Hornauer, S. S. Kumar, M. Schröter, K. Borgwardt, and A. Hierlemann. Scalable covariance-based connectivity inference for synchronous neuronal networks.

[46] D. C. Krakauer. Symmetry–simplicity, broken symmetry–complexity. Interface Focus, 13, 4 2023.

[47] M. L. Kringelbach, Y. S. Perl, E. Tagliazucchi, and G. Deco. Toward naturalistic neuroscience: Mechanisms underlying the flattening of brain hierarchy in movie-watching compared to rest and task. Science Advances, 9, 1 2023.

[48] C. Kwon, P. Ao, and D. J. Thouless. Structure of stochastic dynamics near fixed points. Proceedings of the National Academy of Sciences, 102:13029–13033, 9 2005.

[49] T. W. Lin, A. Das, G. P. Krishnan, M. Bazhenov, and T. J. Sejnowski. Differential covariance: A new class of methods to estimate sparse connectivity from neural recordings. Neural Computation, 29:2581–2632, 10 2017.

[50] A. Liska, A. Galbusera, A. J. Schwarz, and A. Gozzi. Functional connectivity hubs of the mouse brain. NeuroImage, 115:281–291, 7 2015.

[51] R. Liégeois, J. Li, R. Kong, C. Orban, D. V. D. Ville, T. Ge, M. R. Sabuncu, and B. T. T. Yeo. Resting brain dynamics at different timescales capture distinct aspects of human behavior. Nature Communications, 10:2317, 5 2019.

[52] C. W. Lynn, E. J. Cornblath, L. Papadopoulos, M. A. Bertolero, and D. S. Bassett. Broken detailed balance and entropy production in the human brain. Proceedings of the National Academy of Sciences of the United States of America, 118:e2109889118, 11 2021.

[53] D. S. Marcus, M. P. Harms, A. Z. Snyder, M. Jenkinson, J. A. Wilson, M. F. Glasser, D. M. Barch, K. A. Archie, G. C. Burgess, M. Ramaratnam, M. Hodge, W. Horton, R. Herrick, T. Olsen, M. McKay, M. House, M. Hileman, E. Reid, J. Harwell, T. Coalson, J. Schindler, J. S. Elam, S. W. Curtiss, and D. C. V. Essen. Human connectome project informatics: quality control, database services, and data visualization. NeuroImage, 80:202–219, 10 2013.

[54] D. S. Margulies, S. S. Ghosh, A. Goulas, M. Falkiewicz, J. M. Huntenburg, G. Langs, G. Bezgin, S. B. Eickhoff, F. X. Castellanos, M. Petrides, E. Jefferies, and J. Smallwood. Situating the default-mode network along a principal gradient of macroscale cortical organization. Proceedings of the National Academy of Sciences of the United States of America, 113:12574–12579, 11 2016.

[55] N. T. Markov, M. Ercsey-Ravasz, D. C. V. Essen, K. Knoblauch, Z. Toroczkai, and H. Kennedy. Cortical high-density counterstream architectures. Science, 342, 11 2013.

[56] E. Nozari, M. A. Bertolero, J. Stiso, L. Caciagli, E. J. Cornblath, X. He, A. S. Mahadevan, G. J. Pappas, and D. S. Bassett. Is the brain macroscopically linear? a system identification of resting state dynamics. bioRxiv, page 2020.12.21.423856, 1 2021.

[57] S. W. Oh, J. A. Harris, L. Ng, B. Winslow, N. Cain, S. Mihalas, Q. Wang, C. Lau, L. Kuan, M. Henry, M. T. Mortrud, B. Ouellette, T. N. Nguyen, S. A. Sorensen, C. R. Slaughterbeck, W. Wakeman, Y. Li, D. Feng, A. Ho, E. Nicholas, K. E. Hirokawa, P. Bohn, K. M. Joines, H. Peng, M. J. Hawrylycz, J. W. Phillips, J. G. Hohmann, P. Wohnoutka, C. R. Gerfen, C. Koch, A. Bernard, C. Dang, A. R. Jones, and H. Zeng. A mesoscale connectome of the mouse brain. Nature 2014 508:7495, 508:207–214, 4 2014.

[58] L. Parkes, J. Z. Kim, J. Stiso, M. E. Calkins, M. Cieslak, R. E. Gur, R. C. Gur, T. M. Moore, M. Ouellet, D. R. Roalf, R. T. Shinohara, D. H. Wolf, T. D. Satterthwaite, and D. S. Bassett. Asymmetric signaling across the hierarchy of cytoarchitecture within the human connectome. Science Advances, 8, 12 2022.

[59] X. Peng, Q. Liu, C. S. Hubbard, D. Wang, W. Zhu, M. D. Fox, and H. Liu. Robust dynamic brain coactivation states estimated in individuals. Science Advances, 9, 1 2023.

[60] Y. S. Perl, H. Bocaccio, C. Pallavicini, I. Pérez-Ipiña, S. Laureys, H. Laufs, M. Kringelbach, G. Deco, and E. Tagliazucchi. Nonequilibrium brain dynamics as a signature of consciousness. Physical Review E, 104:014411, 7 2021.

[61] A. Ponce-Alvarez and G. Deco. The hopf whole-brain model and its linear approximation. 9 2023.

[62] G. Prando, M. Zorzi, A. Bertoldo, M. Corbetta, M. Zorzi, and A. Chiuso. Sparse dcm for whole-brain effective connectivity from resting-state fmri data. NeuroImage, 208, 2020.

[63] G. Rabuffo, J. Fousek, C. Bernard, and V. Jirsa. Neuronal cascades shape whole-brain functional dynamics at rest. eNeuro, 8, 9 2021.

[64] A. Razi, M. L. Seghier, Y. Zhou, P. McColgan, P. Zeidman, H. J. Park, O. Sporns, G. Rees, and K. J. Friston. Large-scale dcms for resting-state fmri. Network neuroscience (Cambridge, Mass.), 1:222–241, 1 2017.

[65] F. Rocchi, C. Canella, S. Noei, D. Gutierrez-Barragan, L. Coletta, A. Galbusera, A. Stuefer, S. Vassanelli, M. Pasqualetti, G. Iurilli, S. Panzeri, and A. Gozzi. Increased fmri connectivity upon chemogenetic inhibition of the mouse prefrontal cortex. Nature Communications 2022 13:1, 13:1–15, 2 2022.

[66] E. T. Rolls, M. Joliot, and N. Tzourio-Mazoyer. Implementation of a new parcellation of the orbitofrontal cortex in the automated anatomical labeling atlas. NeuroImage, 122:1–5, 11 2015.

[67] S. Ryali, T. Chen, A. Padmanabhan, W. Cai, and V. Menon. Development and validation of consensus clustering-based framework for brain segmentation using resting fmri. Journal of Neuroscience Methods, 240:128–140, 1 2015.

[68] A. Schaefer, R. Kong, E. M. Gordon, T. O. Laumann, X.-N. Zuo, A. J. Holmes, S. B. Eickhoff, and B. T. T. Yeo. Local-global parcellation of the human cerebral cortex from intrinsic functional connectivity mri. Cerebral Cortex, 28:3095–3114, 9 2018.

[69] M. Schirner, X. Kong, B. T. Yeo, G. Deco, and P. Ritter. Dynamic primitives of brain network interaction. NeuroImage, 250:118928, 4 2022.

[70] C. Seguin, A. Razi, and A. Zalesky. Inferring neural signalling directionality from undirected structural connectomes. Nature Communications 2019 10:1, 10:1–13, 9 2019.

[71] M. Shanahan. Metastable chimera states in community-structured oscillator networks. Chaos: An Interdisciplinary Journal of Nonlinear Science, 20, 3 2010.

[72] M. Shinn, A. Hu, L. Turner, S. Noble, K. H. Preller, J. L. Ji, F. Moujaes, S. Achard, D. Scheinost, R. T. Constable, J. H. Krystal, F. X. Vollenweider, D. Lee, A. Anticevic, E. T. Bullmore, and J. D. Murray. Functional brain networks reflect spatial and temporal autocorrelation. Nature Neuroscience 2023 26:5, 26:867–878, 4 2023.

[73] E. Silvestri, M. Moretto, S. Facchini, M. Castellaro, M. Anglani, E. Monai, D. D’Avella, A. D. Puppa, D. Cecchin, A. Bertoldo, and M. Corbetta. Widespread cortical functional disconnection in gliomas: an individual network mapping approach. Brain Communications, 4, 3 2022.

[74] V. Sip, M. Hashemi, T. Dickscheid, K. Amunts, S. Petkoski, and V. Jirsa. Characterization of regional differences in resting-state fmri with a data-driven network model of brain dynamics. Science Advances, 9, 3 2023.

[75] P. H. Siu, E. Müller, V. Zerbi, K. Aquino, and B. D. Fulcher. Extracting dynamical understanding from neural-mass models of mouse cortex. Frontiers in Computational Neuroscience, 16:847336, 4 2022.

[76] P. Sorrentino, G. Rabuffo, F. Baselice, E. T. Lopez, M. Liparoti, M. Quarantelli, G. Sorrentino, C. Bernard, and V. Jirsa. Dynamical interactions reconfigure the gradient of cortical timescales. Network Neuroscience, 7:73–85, 1 2023.

[77] J. Wang. Landscape and flux theory of non-equilibrium dynamical systems with application to biology. Advances in Physics, 64:1–137, 1 2015.

[78] P. Wang, R. Kong, X. Kong, R. Liégeois, C. Orban, G. Deco, M. P. van den Heuvel, and B. T. Yeo. Inversion of a large-scale circuit model reveals a cortical hierarchy in the dynamic resting human brain. Science Advances, 5, 1 2019.

[79] Q. Wang, S.-L. Ding, Y. Li, J. Royall, D. Feng, P. Lesnar, N. Graddis, M. Naeemi, B. Facer, A. Ho, T. Dolbeare, B. Blanchard, N. Dee, W. Wakeman, K. E. Hirokawa, A. Szafer, S. M. Sunkin, S. W. Oh, A. Bernard, J. W. Phillips, M. Hawrylycz, C. Koch, H. Zeng, J. A. Harris, and L. Ng. The allen mouse brain common coordinate framework: A 3d reference atlas. Cell, 181:936–953.e20, 5 2020.

[80] Y. Xu, X. Long, J. Feng, and P. Gong. Interacting spiral wave patterns underlie complex brain dynamics and are related to cognitive processing. Nature Human Behaviour 2023 7:7, 7:1196–1215, 6 2023.

[81] D. Xue. Conservation-dissipation structure of linear stochastic systems. pages 5980–5985. IEEE, 12 2016.

[82] R. K. P. Zia and B. Schmittmann. Probability currents as principal characteristics in the statitical mechanics of non-equilibrium steady states. Journal of Statistical Mechanics: Theory and Experiment, 2007:P07012–P07012, 7 2007.

